# Compellingly high SARS-CoV-2 susceptibility of Golden Syrian hamsters suggests multiple zoonotic infections of pet hamsters during the COVID-19 pandemic

**DOI:** 10.1101/2022.04.19.488826

**Authors:** Claudia Blaurock, Angele Breithaupt, Saskia Weber, Claudia Wylezich, Markus Keller, Björn-Patrick Mohl, Dirk Görlich, Martin H. Groschup, Balal Sadeghi, Dirk Höper, Thomas C. Mettenleiter, Anne Balkema-Buschmann

## Abstract

Golden Syrian hamsters (*Mesocricetus auratus*) are used as a research model for severe acute respiratory syndrome coronavirus type 2 (SARS-CoV-2). Millions of Golden Syrian hamsters are also kept as pets in close contact to humans. To determine the minimum infective dose (MID) for assessing the zoonotic transmission risk, and to define the optimal infection dose for experimental studies, we orotracheally inoculated hamsters with SARS-CoV-2 doses from 1*10^5^ to 1*10^−4^ tissue culture infectious dose 50 (TCID_50_). Body weight and virus shedding were monitored daily. 1*10^−3^ TCID_50_ was defined as the MID, and this was still sufficient to induce virus shedding at levels up to 10^2.75^ TCID_50_/ml, equaling the estimated MID for humans. Virological and histological data revealed 1*10^2^ TCID_50_ as the optimal dose for experimental infections. This compellingly high susceptibility resulting in productive infections in Golden Syrian hamsters needs to be considered also as a source of SARS-CoV-2 infections in humans.

## Introduction

To date, more than 499 million confirmed Coronavirus Disease 2019 (COVID-19) cases and more than 6.1 million deaths have been reported globally since the initial detection of severe acute respiratory syndrome coronavirus 2 (SARS-CoV-2) in December 2019 (https://coronavirus.jhu.edu; 12.04.2022). COVID-19 has been studied extensively since, and the incubation time in humans has been calculated as 6.4 days for the initially described virus variants (Cheng et al., 2021; Elias, Sekri, Leblanc, Cucherat, & Vanhems, 2021), while the estimated minimum infective dose (MID) for humans has been calculated to be approximately 10^2^ tissue culture infectious dose 50 (TCID_50_) (Basu, 2021; Karimzadeh, Bhopal, & Nguyen Tien, 2021).

Golden Syrian hamsters (*Mesocricetus auratus*) are widely used as a COVID-19 animal model, as they efficiently support SARS-CoV-2 replication and display pathological manifestations similar to human COVID-19 pneumonia, such as focal diffuse alveolar destruction, hyaline membrane formation, and mononuclear cell infiltration (Chan et al., 2020; Sia et al., 2020). Efficient transmission from inoculated to naïve hamsters by direct contact and aerosol has also been reported (Dowall et al., 2021; Sia et al., 2020). Several hamster species are kept as pets in millions of households worldwide. These include Golden Syrian hamsters, Chinese hamsters (*Cricetulus griseus*), and three Dwarf hamster species (*Phodopus roborowskii, P. campbelli* and *P. sungorus*). According to the German Pet Trade & Industry Association (Zentralverband Zoologischer Fachbetriebe e.V. ZZF), approximately 520.000 pet hamsters were kept in German households in 2020. Golden Syrian hamsters comprised 43% (223.600 animals), and the Dwarf hamster species 54% (280.800 animals) (source: Zentralverband Zoologischer Fachbetriebe (ZZF) and Industrieverband Heimtierbedarf (IVH)). While Golden Syrian hamsters, Chinese hamsters (*Cricetulus griseus*), and Roborovski Dwarf hamsters (*Phodopus roborovskii*) are highly susceptible and develop clinical disease and lung pathology, both other Dwarf hamster species (Campbell’s Dwarf hamster and Djungarian Dwarf hamster) only developed subclinical disease with mild tissue damage and a rapid onset of regeneration under experimental conditions (Bertzbach et al., 2021; Trimpert et al., 2020).

Since the onset of the pandemic, several virus variants have emerged, five of which have been classified as variants of concern (VOC) by the World Health Organization (WHO) for their enhanced transmissibility and virulence or their reduced susceptibility to immune reactions (WHO, 01.03.2022). These include variants B.1.1.7 (Alpha), B.1.351 (Beta), P.1 (Gamma), B.1.617.2 (Delta) and B.1.1.529 (Omicron). Especially the rapid worldwide spread of the Delta and Omicron variants have raised concern about an acceleration of the pandemic’s dynamic, despite increasing proportions of fully vaccinated humans (https://github.com/owid/covid-19-data/tree/master/public/data/vaccinations). Therefore, the pathogenesis induced by VOCs has been addressed in a number of recent animal studies, mostly performed in Golden Syrian hamsters (Abdelnabi et al., 2022; McMahan et al., 2022; Sreelekshmy Mohandas et al., 2022; S. Mohandas et al., 2021; O’Donnell et al., 2021). While Alpha, Beta and Delta variants caused clinical manifestation and pathology similar to the ancestral virus (S. Mohandas et al., 2021; O’Donnell et al., 2021), infection with the Omicron variant caused significantly milder clinical signs and lower levels of pulmonary affection (Abdelnabi et al., 2022; McMahan et al., 2022). However, levels of viral genomic RNA, as well as subgenomic RNA (sgRNA), were distinctly higher in the nasal turbinate samples, indicating a higher transmission risk from these animals as compared to animals infected with any of the other variants (McMahan et al., 2022).

Very recently, two distinct zoonotic transmission events from hamsters to humans were reported from Hong Kong followed by human-to-human transmission (Yen, 2022), after the import of pet Golden Syrian hamsters from The Netherlands, causing a re-introduction of the Delta VOC into the country. Investigation of these animals revealed SARS-CoV-2 RNA in 15 out of 28 Golden Syrian hamsters, but in none of the 77 analysed Dwarf hamsters (Yen, 2022). These findings gave rise to serious concern about the SARS-CoV-2 transmission risk from pet hamsters to humans in affected households (Haagmans & Koopmans, 2022; Kok et al., 2022), and more than 2.000 pet hamsters were subsequently culled in Hong Kong. This highlights the importance of an evidence-based risk assessment regarding SARS-CoV-2 transmissions from pet hamsters to humans and vice versa. To the best of our knowledge, no systematic quantitative study on the susceptibility of Golden Syrian hamsters to a SARS-CoV-2 infection has been undertaken so far. We therefore infected groups of Golden Syrian hamsters with serial SARS-CoV-2 dilutions from 10^5^ to 10^−4^ TCID_50_, which is equivalent to 0.7 genome copy numbers per dose, as determined by reverse transcriptase quantitative real-time PCR of serial dilutions of a SARS-CoV-2 RNA standard. Animals infected with a dose of 1*10^−3^ TCID_50_ shed replication competent virus, and accumulated SARS-CoV-2 infectivity in their respiratory tract until 7 days post inoculation (dpi). This study demonstrates the extremely high SARS-CoV-2 susceptibility of Golden Syrian hamsters. For drug or vaccine efficacy studies, a moderate infection dose, sufficiently high to induce reproducible viral loads and histological findings, and at the same time sufficiently low to avoid a highly artificial infection should be applied, which in our experimental setup was determined as 1*10^2^ TCID_50_.

## Results

### Pronounced weight loss and high levels of viral shedding upon orotracheal inoculation

Upon orotracheal and intranasal inoculation with SARS-CoV-2 (Fig. 1A), animals started losing body weight from 2 dpi, and the mean weight loss at 7 dpi was 16% after orotracheal inoculation and 7% after intranasal inoculation (Fig. 1 B). We observed a statistically significant 5-to 10-fold increase in viral genome shedding in the nasal wash samples at 6 and 9 dpi after orotracheal inoculation as compared to the intranasal inoculation (Fig. 1C). Upon necropsy at 14 dpi, viral RNA in the respiratory tract samples was not detectable at levels allowing a quantitative comparison, and no replication competent virus was detected. Based on these data, we decided to continue with the orotracheal inoculation for subsequent experiments.

**Figure 1:**
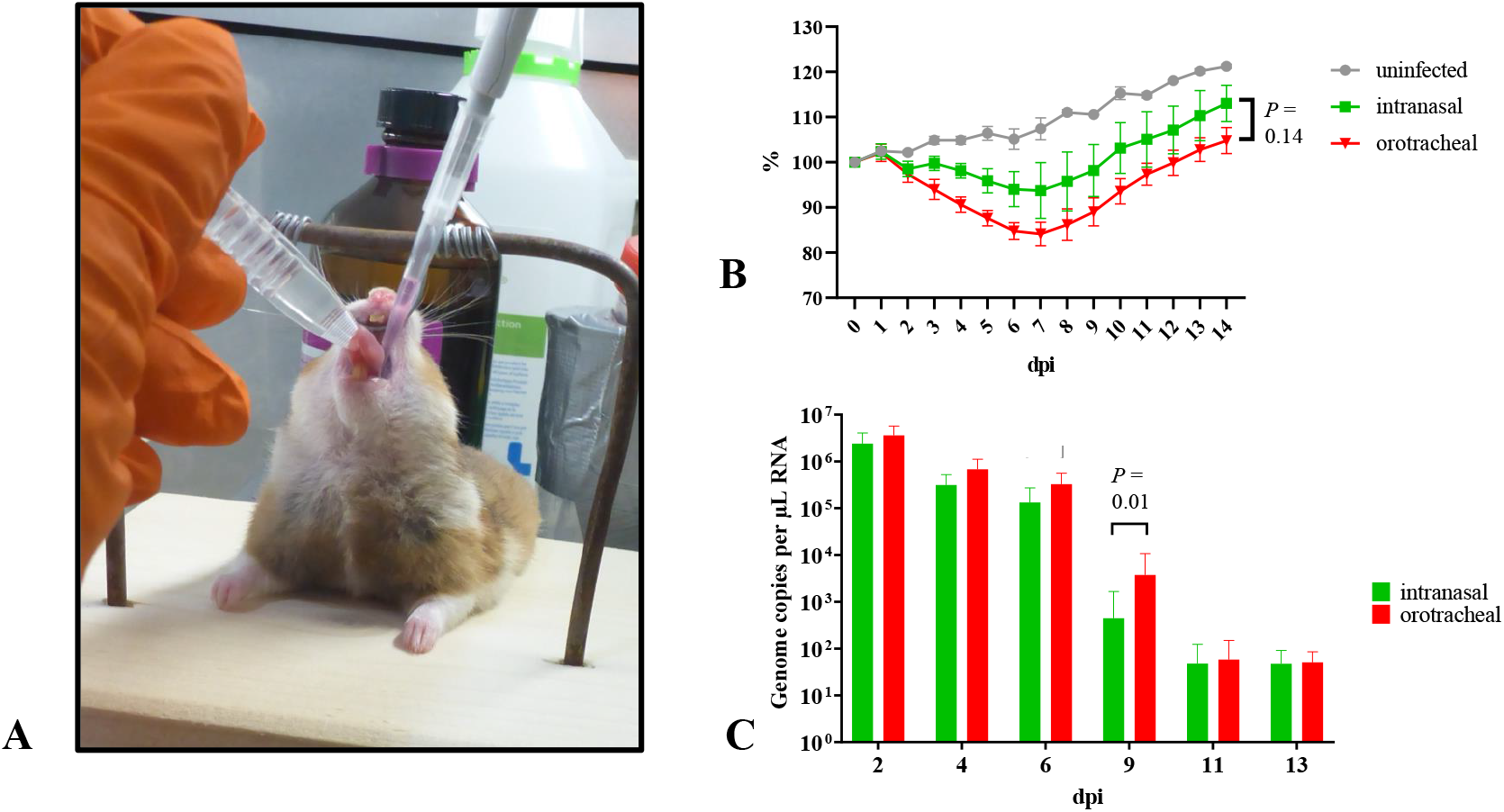
Orotracheal SARS-CoV-2 inoculation results in increased weight loss and viral shedding. (A) Inoculation technique. Anaesthetized animals are fixed with a stretched neck and extended tongue, inoculum is instilled on the root of the tongue. (B) Body weight curves of uninfected control group (grey), groups infected by the orotracheal (red) and nasal routes (green); *N* = 19. (C) Oral and nasal shedding between 1 and 13 dpi after orotracheal (red) and intranasal (green) inoculation; *N* = 16.

### 1*10^−1^ TCID_50_ SARS-CoV-2 is sufficient to induce body weight reduction and pneumonia

Animals inoculated in our first study with doses exceeding 1*10^2^ TCID_50_ SARS-CoV-2 showed a decrease in body weight from 2 dpi, while hamsters inoculated with 1*10^1^ TCID_50_ gained weight until 5 dpi, before losing weight until the end of the experiment. Animals inoculated with 1*10^−1^ and 1*10^0^ TCID_50_ as well as the uninfected controls continued to gain weight throughout the experiment. In the second study, all groups receiving 1*10^−1^ TCID_50_ or higher lost weight until the end of the experiment, with an onset delayed by 2-3 day in animals infected with 1*10^1^ and 1*10^0^ TCID_50_. In a third study using dilutions to 1*10^−4^ TCID_50_ and continuing until 10 dpi, doses of 10^−1^ TCID_50_ or higher induced weight loss (Fig. 2). Statistical analysis revealed significant differences between the body weights determined per group at the day of the necropsy, i.e. 7 and 10 dpi (Suppl. Tab. 1).

**Figure 2:**
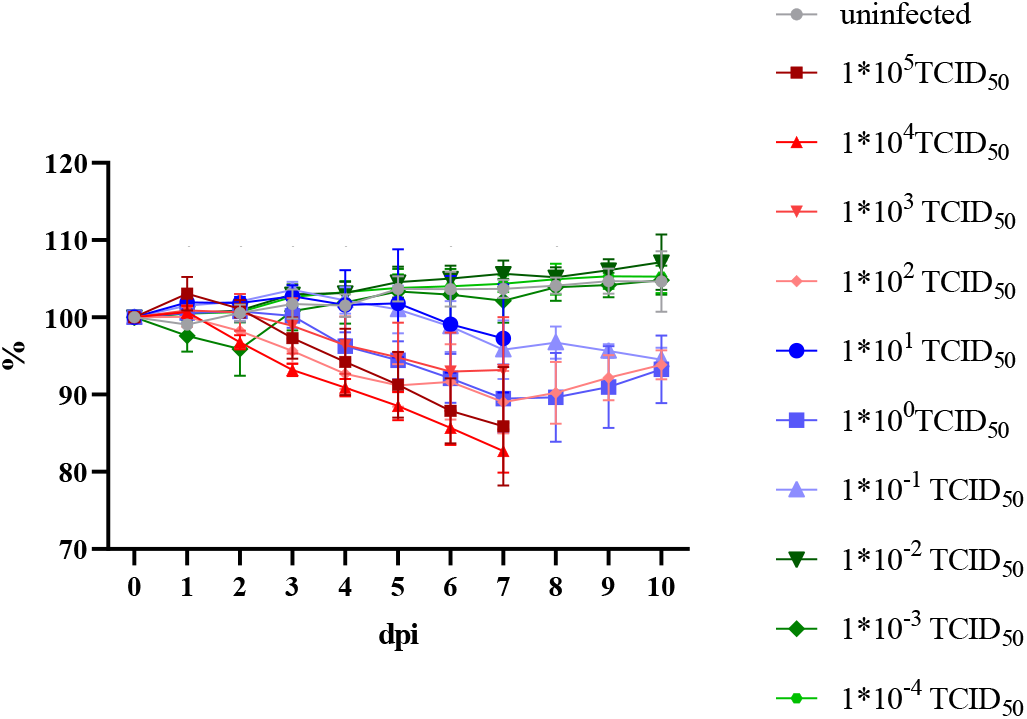
Mean and standard deviation of body weight per infection dose group. Changes in body weight (%) in relation to 0 dpi. Statistically significant differences in daily body weight changes are marked by an asterisk. *N* = 69, *P* = 0.0001 for all days. Further details on statistical analysis are shown in Suppl. Tab. 1.

During autopsy performed at 7 or 10 dpi, the extent of reddish discoloration of the whole lung was recorded. We observed decreasing estimated levels of lung affection correlated with decreasing infection doses. We still noted areas of discoloration at 7 dpi in animals that had been infected with 1*10^−3^ TCID_50_ (Fig.3 A). At 10 dpi, even hamsters inoculated with the lowest dose of 1*10^−4^ TCID_50_ showed lung changes (Fig. 3 B).

**Figure 3:**
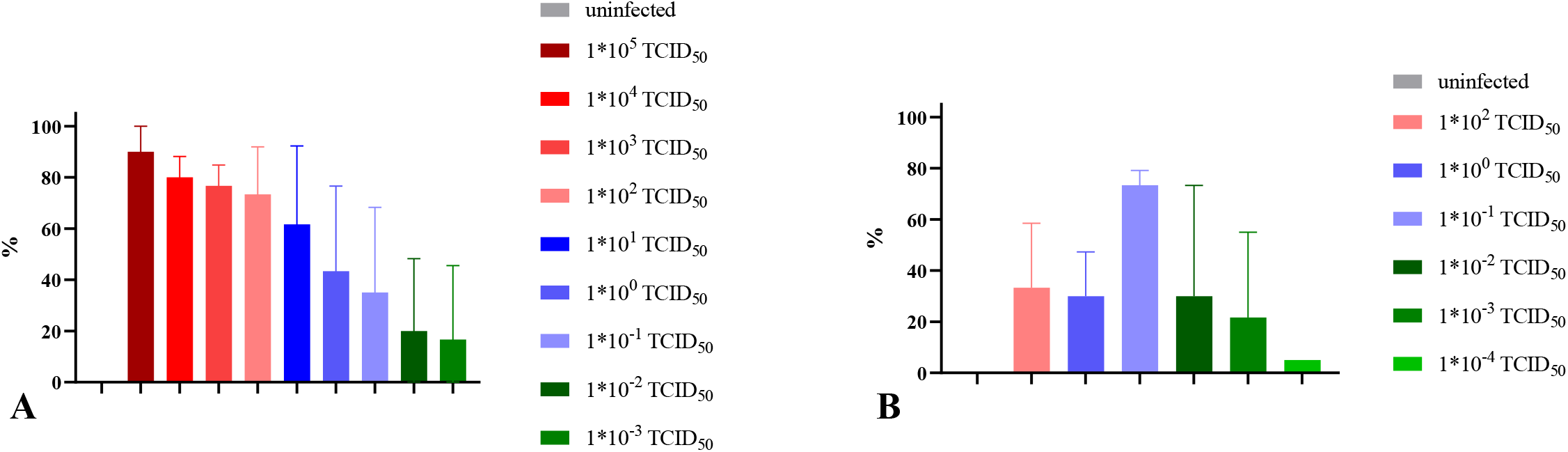
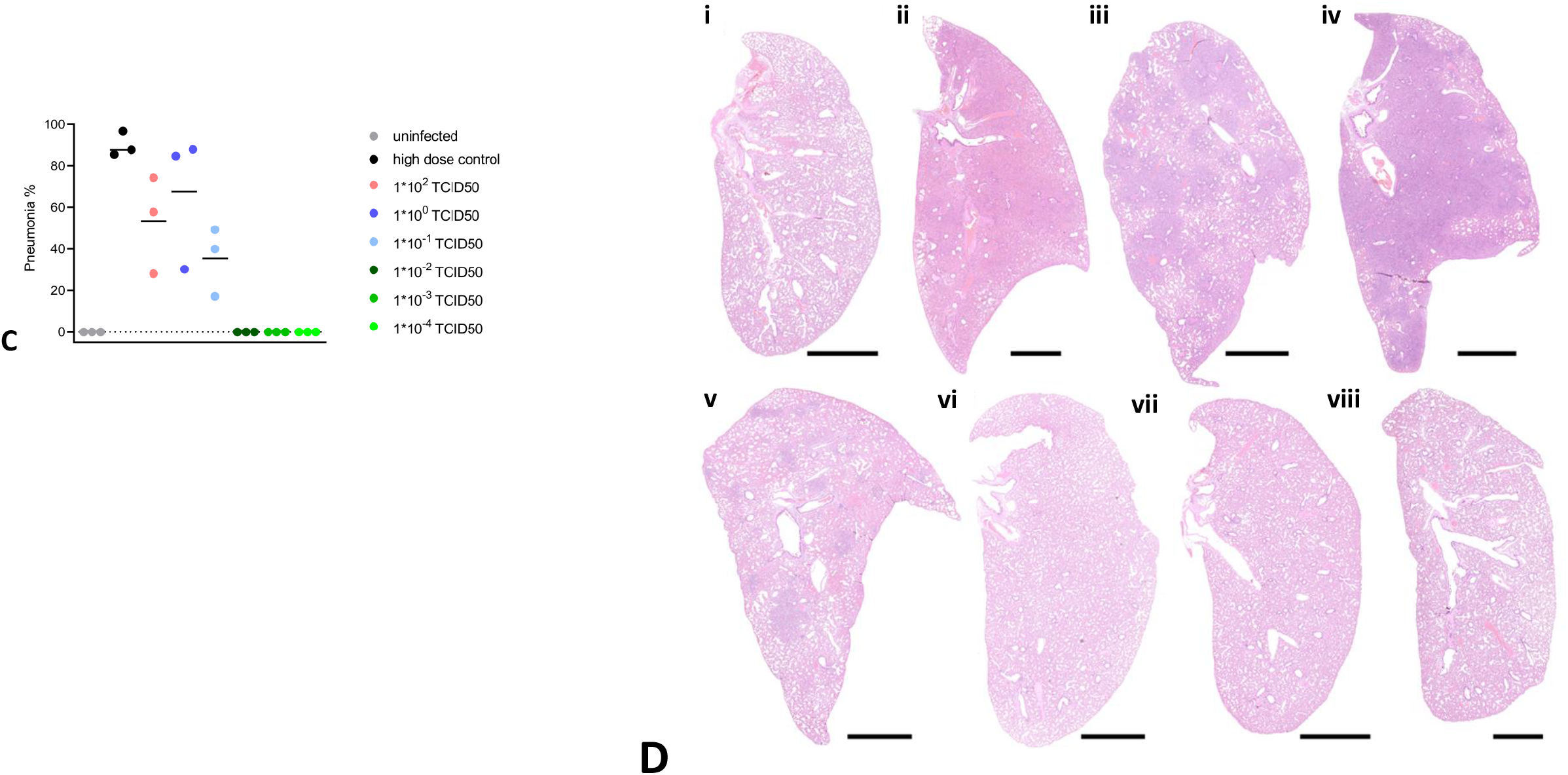
Levels of lung affection after challenge with different infection doses. (A) macroscopically assessed level of affection of whole lung (%) during autopsy at 7 dpi and (B) at 10 dpi displayed as the mean value and standard deviation for each group. (C-D) Histopathology of hamster lungs, 10 days after orotracheal SARS-CoV-2 infection. Pneumonia-associated consolidation (C) and representative overviews (D) of the entire left lung lobe of (i) uninfected control; (ii) high dose control; (iii) infected with 1*10^2^ TCID_50_; (iv) infected with 1*10^0^ TCID_50_; (v) infected with 1*10^−1^ TCID_50_; (vi) infected with 1*10^−2^ TCID_50_; (vii) infected with 1*10^−3^ TCID_50_; (viii) infected with 1*10^−4^ TCID_50_. (B-D) n = 3.

Histopathology was performed on the left lung lobe of animals sacrificed at 10 dpi. Archived lungs from a high-dose infection study (1*10^4.5^ TCID_50_) served as positive controls. Pneumonia-associated consolidation was consistently found in all hamsters infected with 1*10^−1^ TCID_50_ SARS-CoV-2 or higher (Fig. 3 C, D). Lungs collected at 10 dpi mainly showed regenerative changes with bronchial and type-2-pneumoncyte hyperplasia and hypertrophy with formation of multinucleated, atypical cells. However, still a moderate to severe inflammatory reaction was found, with intra-alveolar, interstitial, peribronchial and perivascular inflammatory infiltrates, as well as vasculitis and/or endothelialitis. Acute necrosis of the bronchial epithelium was a rare finding (Fig. 4 A - C). Although not statistically significant, hamsters infected with 1*10^−1^ TCID_50_ showed a tendency to be less severely affected (Fig. 4 E). In general, animals infected with doses of 1*10^−2^ TCID_50_ and lower did not develop pneumonia-associated consolidation in the lung (Fig. 4 F, G). However, focal alveolar or perivascular infiltrates or bronchial epithelial hyperplasia were found in individual animals in each group indicating prior local infection, even after infection with 1*10^−4^ TCID_50_ (Fig. 4 H, I).

**Figure 4:**
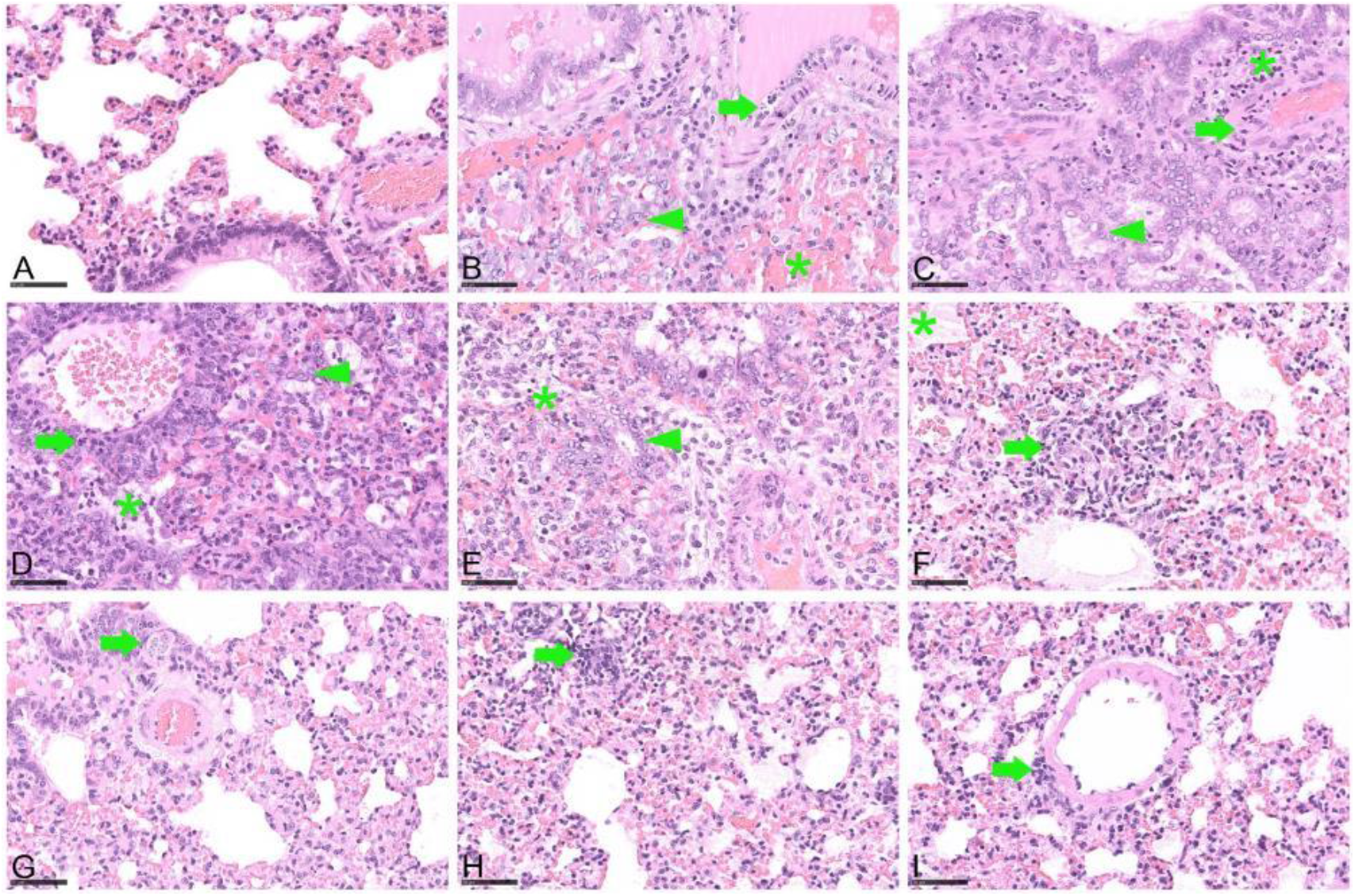
Detailed histopathology of hamster lungs, 10 days after orotracheal SARS-CoV-2 infection. Hematoxylin and eosin staining, bar 50 µm. (i) Uninfected control, no lung lesion, (ii) after 1*10^4.5^ TCID_50_ infection, showing vasculitis (arrow), type 2 pneumocyte hyperplasia and bronchialisation of alveoli (arrowhead), intra-alveolar erythrocytes (asterisk), (iii) after 1*10^2^ TCID_50_ infection, with vasculitis (arrow), type 2 pneumocyte hyperplasia and bronchialisation of alveoli (arrowhead) and perivascular infiltrates (asterisk), (iv) after 1*10^0^ TCID_50_ infection exhibiting perivascular (arrow) and alveolar (asterisk) infiltrates, type 2 pneumocyte hyperplasia and bronchialisation of alveoli (arrowhead), (v) after 1*10^−1^ TCID_50_ infection, showing alveolar infiltrates (asterisk), type 2 pneumocyte hyperplasia and bronchialisation of alveoli (arrowhead), (vi) after 1*10^−2^ TCID_50_ infection, with focal alveolar infiltrates (arrow) and edema (asterisk), (vii) after 1*10^−3^ TCID_50_ infection, exhibiting focal hypertrophy/hyperplasia of bronchial epithelium (arrow), (viii) after 1*10^−4^ TCID_50_, with focal alveolar infiltrates (arrow), (ix) after 1*10^−4^ TCID_50_ showing focal perivascular infiltrates (arrow).

### Lateral flow device Rapid Test detects infections with 1*10^−1^ TCID_50_ SARS-CoV-2

We analysed oral swab samples collected at 6 dpi using the Nowcheck COVID-19 Rapid Test, which confirmed the infection in hamsters inoculated with doses higher than 1*10^0^ TCID_50_, as well as in two hamsters inoculated with 1*10^−1^ TCID_50_, which translates into samples with ct-values in the real-time RT-PCR below 30. All uninfected hamsters as well as one hamster inoculated with 1*10^−1^ TCID_50_ were negative at 6 dpi (Fig. S1). We also analysed nasal washes collected at 7 dpi Again, the infection was detectable in animals challenged with doses of 1*10^−1^ TCID_50_ or higher, i.e. for samples with ct-values below 30.

### 1*10^−2^ TCID_50_ SARS-CoV-2 is sufficient to induce seroconversion

To confirm a systemic infection, all sera collected during necropsies were tested in an indirect ELISA based on the SARS-CoV-2 RBD antigen. ROC analysis was performed for the determination of the ELISA cut-off value. We determined a specificity of 100% and a sensitivity of 99.56% using the cut-off value of 14.11 percent of the positive control value (PP) (Fig. S2 A). We detected seroconversion with a dose-dependent increase of PP values in hamsters inoculated with infection doses of 1*10^−2^ TCID_50_ or higher (Fig. S2 B).

### 1*10^−3^ TCID_50_ SARS-CoV-2 is sufficient to induce oral and nasal viral shedding

From the first day post infection, animals infected with 10^2^ TCID_50_ SARS-CoV-2 or higher shed virus between 10^3.5^ and 10^4.5^ genome copy numbers / µl RNA. By 3 dpi, all animals infected with doses of 1*10^−1^ TCID_50_ or higher shed virus (Fig. 5 A). The oral swab samples contained replicating virus even in the groups infected with a dose of 10^−1^ TCID_50_ until 6 dpi, and at a level of 10^2^ TCID_50_ / ml in the swab sample collected at 6 dpi from one animal infected with the 1*10^−3^ TCID_50_ dose. This result was confirmed by the detection of SARS-CoV-2 sgRNA as an additional proof of a present or past virus replication. All positive results for the nasal wash samples collected at 2 and 4 dpi were confirmed by re-testing for sgRNA, with ct values of about 3 points above those determined for the SARS-CoV-2 N-gene (Tab. 2). The peak of oral shedding of replication competent virus was observed between 3 and 5 dpi, at levels reaching up to 10^3^ TCID_50_ / ml in the oral swab samples (Fig 5 B).

**Figure 5:**
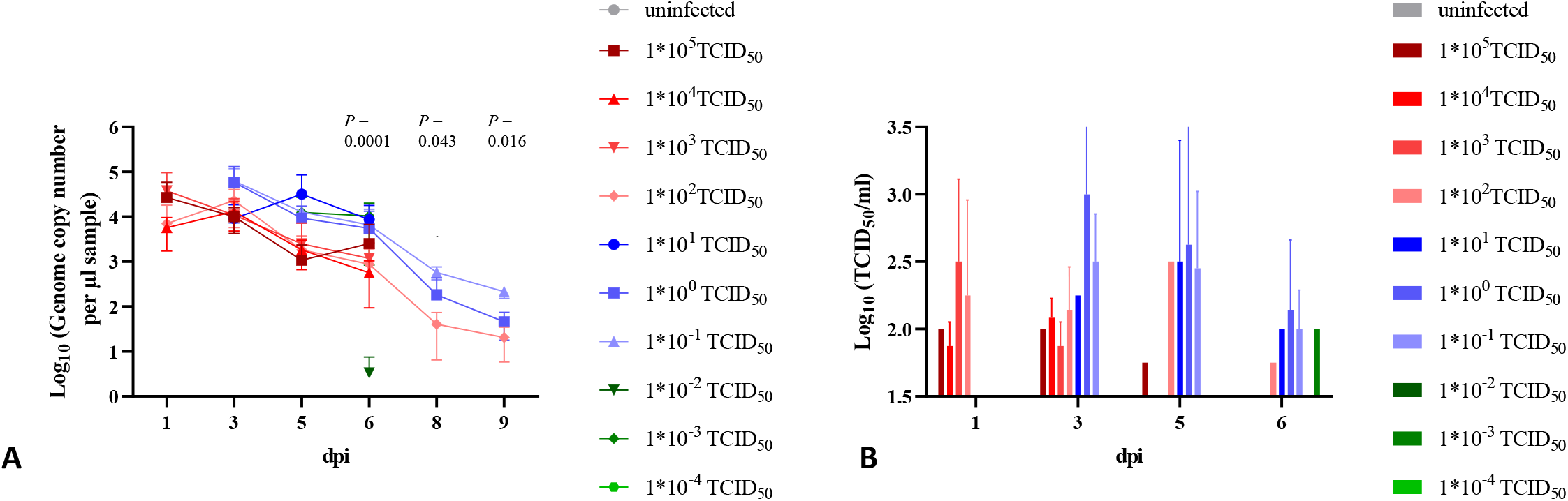
Viral shedding in oral swab samples. (A) Genome copy number (Log10) of viral RNA and (B) replication competent virus (Log10 TCID_50_/ml) detected in the oral swab samples of all hamsters. All groups were analysed, negative results are not shown. p-values are indicated for statistically significant differences in RNA levels. Details on statistical analysis are shown in Suppl. Tab. 1. Colors match and represent results from the same hamsters as shown before and following.

SARS-CoV-2 RNA levels detected in the nasal washes were generally 10-to100-times higher than those determined from the oral swab samples. We were able to detect viral RNA at 2 and 4 dpi in nasal washes, even in the animals infected with the lowest dose of 10^−4^ TCID_50_ (Fig. 6 A). Significant differences were determined for the copy numbers detected in the different infection groups (Tab 1). The peak of nasal shedding of replication competent virus was observed at 2 dpi at levels reaching up to 10^7^ TCID_50_ / ml, while 1*10^2.75^ TCID_50_ (equivalent to the estimated MID for humans), as well as sgRNA were detected in a sample collected from one animal infected with 10^−3^ TCID_50_ (Fig. 6 B). The dose-dependent delay in onset of oral and nasal shedding is summarized in Fig. 7.

**Figure 6:**
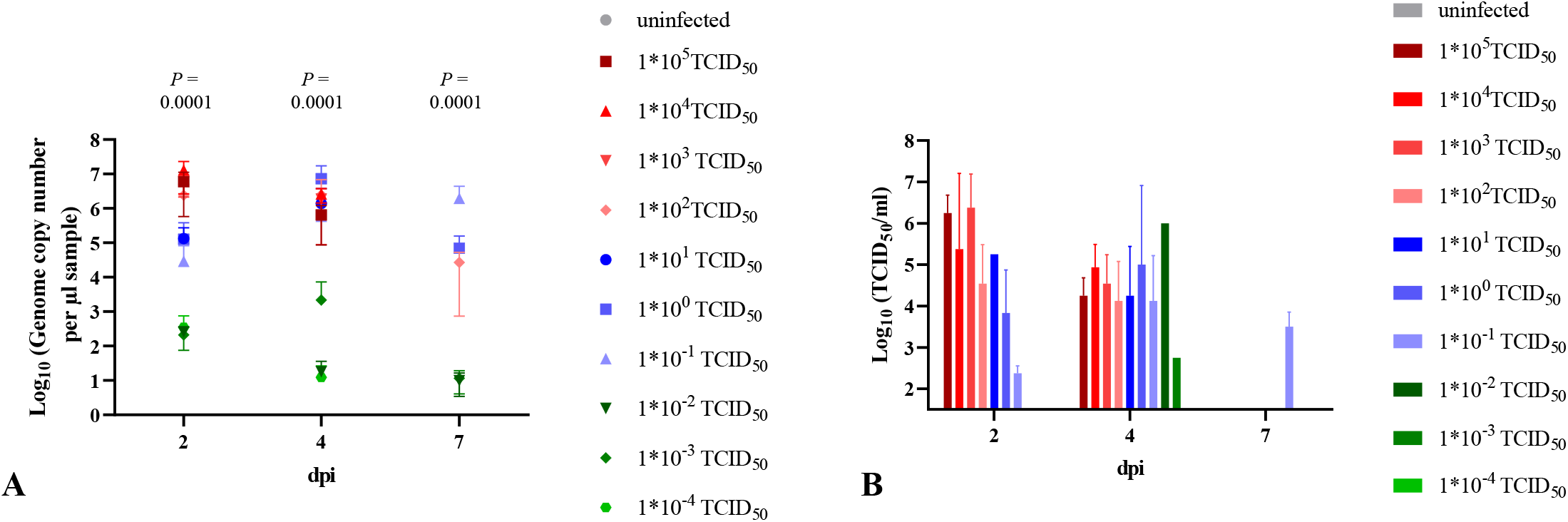
Viral shedding in nasal washes. (A) Genome copy number (Log10) of SARS-CoV-2 RNA and (B) replication competent virus (Log10 TCID_50_/ml) detected in the nasal wash samples of all hamsters. All groups were analysed, negative results are not shown. Details on statistical analysis are shown in Suppl. Tab. 1. Colors match and represent results from the same hamsters as shown before and following.

**Figure 7:**
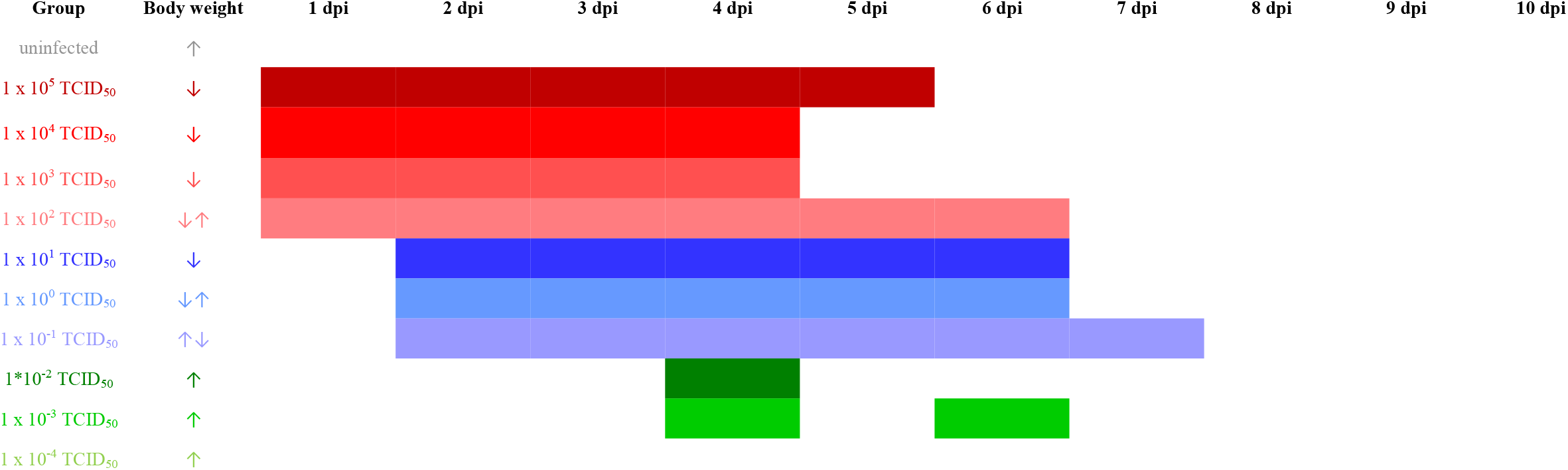
Time-dependent shedding of replication competent virus and change in body weight after infection with decreasing SARS-CoV-2 doses. Body weight development is shown by arrows; ↑ increasing from 1-7 or 10 dpi; ↓ decreasing from 1 – 7 or 10 dpi; ↓↑ decrease until 7 dpi followed by increase; ↑↓ increase until 4 dpi, followed by decrease until 10 dpi. Colors match and represent results from the same hamsters as shown before and following.

### Replication competent virus in lungs after infection with 1*10^−1^ TCID_50_, and in nasal conchae after infection with 1*10^−3^ TCID_50_

Similar levels of viral RNA were detected in nasal conchae samples of all animals sacrificed at 7 dpi, independent of the inoculation dose, including the group infected with a dose of 1*10^−3^ TCID_50_ (Fig. 8 A), while at 10 dpi, viral RNA was only detected in the samples collected from the infection groups 1*10^2^ to 1*10^−1^ TCID_50_ (Fig. 8 B). Interestingly at 7 dpi, the highest levels of viral RNA were detected in trachea and lung samples collected in groups inoculated with the lowest infection dose of 1*10^−3^ TCID_50_. At this time point, the highest virus titers were determined for the nasal conchae samples collected from anima8ls infected with SARS-CoV-2 doses between 10^0^ and 10^−3^ TCID_50_ (Fig. 8 C), which was confirmed by the detection of sgRNA (Tab. 2), while only nasal conchae samples from animals infected with 10^−1^ TCID_50_ contained detectable levels of replication competent virus at 10 dpi (Fig. 8 D). Statistically different levels of replication competent virus were detected in lung samples collected from animals inoculated with different infection doses (Suppl. Tab. 1).

**Figure 8:**
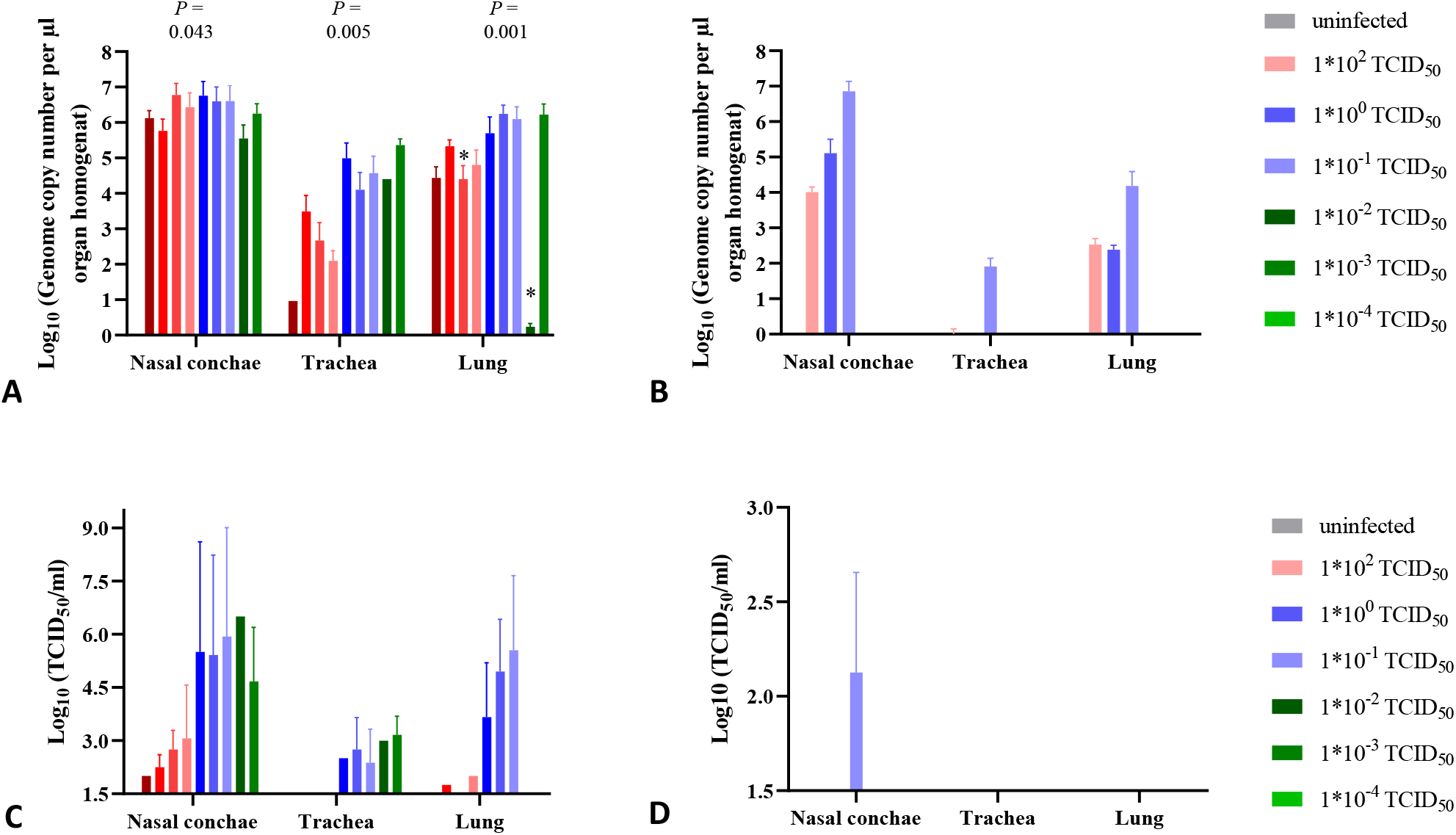
Genome copy number (Log10) of SARS-CoV-2 RNA and replication competent virus (Log10 TCID50/ml) detected in the respiratory tract. (A) Genome copy number (Log10) of viral RNA and (B) replication competent virus (Log10 TCID_50_/ml) detected in the respiratory tract samples at 7 dpi. (C) Genome copy number (Log10) of viral RNA and (D) replication competent virus (Log10 TCID_50_/ml) detected in the respiratory tract samples at 10 dpi. Statistically significant differences in RNA levels were determined at 7 dpi. Further details on statistical analysis are shown in Suppl. Tab. 1 and Suppl. Tab. 3. Colors match and represent results from the same hamsters as shown before.

### Sequence analysis of SARS-CoV-2 retrieved from animals infected with a high or a low dose

We exemplarily analysed SARS-CoV-2 whole-genome sequences retrieved from nasal conchae samples of two animals. Animal H310 (L5497) from a high-dose infection study (1*10^4,5^ TCID_50_) autopsied at 10 dpi, whereas animal H318 (L5537) was infected with 1*10^−1^ TCID_50_. Both animals and was sacrificed at 10 dpi. The sequence of the first sample is identical to the virus stock 2019_nCoV Muc-IMB-1 and shows only one SNV, a transition in the M gene (C26877T) with 52% variant frequency (strand bias 51%) that was already detected as minor variant with 18% variant frequency in the virus stock. The sequence of the second animal is identical to the virus stock and shows only one transition in the ORF1a, nsp6 (T11639C) and no further SNVs. Both described changes are silent mutations. The genome sequences are available under ENA study accession number PRJEB51977.

## Discussion

Today, Golden Syrian hamsters are the leading animal model in SARS-CoV-2 research, in particular for vaccine and drug efficacy studies (Chiba et al., 2022; Meseda et al., 2021; Taylor et al., 2021; Yadav et al.), since they mirror the moderate disease phenotype of human patients with a complete recovery within 14 days (Gruber, Firsching, Trimpert, & Dietert, 2021). When establishing this model, we initially used an infection dose of 1*10^5^ TCID_50_, and found the orotracheal infection route to induce a more prominent weight loss and viral shedding, the latter exceeding those obtained after intranasal infection by at least a factor of 5. Therefore, this route was used for all subsequent studies. Since the intranasal route was used in almost all other published studies (Chiba et al., 2022; Meseda et al., 2021; Taylor et al., 2021; Yadav et al.), and to the best of our knowledge no report of an orotracheal SARS-CoV-2 inoculation of hamsters has been published so far, a direct comparison between previously published experiments and our study is not feasible.

In our study, we were able to show that an infection dose of 1*10^−1^ TCID_50_ was sufficient to induce a significant weight loss, oral and nasal shedding as well as accumulation of replication competent virus in the respiratory tract, and SARS-CoV-2 related pneumonia. In this group, the onset of weight loss and lethargy were delayed by 3 days, presumably due to an initial virus replication at the inoculation site, before spreading throughout the respiratory tract. Infection doses of 1*10^−2^ TCID_50_ or higher were sufficient to induce a dose-dependent seroconversion within 7 days, therefore the virus must have reached the lymphatic system within the first 2-3 days post infection. As no blood samples were available for time points before 7 dpi, a possible viremia within the first 3 days post infection could not be monitored. The estimated proportion of affected lung tissue also showed a clear correlation to the infection dose. Even after challenge with 1*10^−3^ TCID_50_, we observed macroscopically visible lung affections, oral and nasal shedding as well as accumulation of replication competent virus in the nasal conchae and the trachea. We therefore define the MID for Golden Syrian hamsters upon orotracheal infection as 1*10^−3^ TCID_50_. This result not only proves the exceptionally high susceptibility of Golden Syrian hamsters to a SARS-CoV-2 infection, but also indicates the low sensitivity of the Vero E6 virus titration assay with a detection limit of 1*10^1.5^ TCID_50_, as compared to 1*10^−3^ TCID_50_ in the Golden Syrian hamster model. In another Golden Syrian hamster susceptibility study published by Rosenke and co-workers (Rosenke et al., 2020) using intranasal infection doses between 1*10^3^ and 1*10^0^ TCID_50_, a dose of 5 TCID_50_ was postulated as MID. The data obtained for the lower dose group resembles what we observed for the group inoculated with 1*10^−2^ TCID_50_, which may be due to the increased efficiency of the orotracheal route. Moreover, since data were only collected until 5 dpi, a delayed onset of virus shedding and virus propagation in the respiratory tract after challenge with very low doses would have remained unnoticed.

We propose an optimal SARS-CoV-2 infection dose for drug or vaccine efficacy studies in Golden Syrian hamsters of 10^2^ TCID_50_, as this is sufficient to induce a reproducible (as confirmed by three independent experiments) weight loss, macroscopically visible lung affection, pneumonia, shedding of replication competent virus from 1 dpi, and the presence of high levels of replication competent virus in the respiratory tract on day 7 post infection, which will therefore allow a quantification of therapeutic effects. Infection doses of 10^5^ TCID_50_ that are generally used in Golden Syrian hamsters (Chan et al., 2020; Dowall et al., 2021; Francis et al., 2021; Johnson et al., 2022) may be too high to allow a disease progression at least partly resembling the distinctly slower processes in humans after a natural infection (Karimzadeh et al., 2021).

Replication competent virus as well as SARS-CoV-2 sgRNA were detectable at dose dependent levels in nasal washes of all groups until 4 dpi except the group infected with 1*10^−4^ TCID_50_. Thus, by this time, the virus must have already disseminated from the trachea and lung (inoculation site) to the upper respiratory tract, which may have occurred either via mucus transport from the lower respiratory tract, or via the bloodstream. Since we did not determine seroconversion in the animals receiving an infection dose of 1*10^−2^ TCID_50_ or lower, virus dissemination via active mucus transport seems most plausible. The positive sgRNA results also indicate a past active viral replication in the nasal epithelium, as sgRNAs are transcribed only in infected cells (Wolfel et al., 2020). However, positive sgRNA results cannot be interpreted as an equivalent to the detection of replication competent virus by TCID_50_ (Alexandersen, Chamings, & Bhatta, 2020).

To allow a first insight whether the exposition to immune reactions in an animal inoculated with a low dose may stimulate the formation of gene mutations, we analysed the SARS-CoV-2 sequences retrieved from nasal conchae samples from one animal each infected with a high dose and a low dose, and found no indication for such an effect.

The confirmation of SARS-CoV-2 infections in animals infected with 1*10^−1^ TCID_50_ or higher by LFD confirms the suitability of these assays for the screening of hamster swab samples, which may be relevant in households with COVID-19 patients where Golden Syrian hamsters are kept as pets. Upon natural infection with a low dose, these animals will most probably not develop clinical signs, but will shed replication competent virus for at least six days at levels possibly exceeding the estimated human MID of 1*10^2^ TCID_50_, thus allowing a zoonotic infection cycle between humans and Syrian Gold hamsters, as it has been observed very recently in Hong Kong (Yen, 2022).

Our data demonstrate the exceptionally high susceptibility of Golden Syrian hamsters to an infection with a German SARS-CoV-2 isolate obtained during an outbreak in Munich in January 2020 (Wolfel et al., 2020). Other groups reported an almost identical disease progression and virus shedding pattern for VOCs Alpha, Beta, and Delta as for the ancestral virus in hamsters (S. Mohandas et al., 2021; O’Donnell et al., 2021), rendering the titration experiments with these VOCs dispensable. Meanwhile, a low pathogenicity combined with high up to 100-fold increased viral loads in the upper respiratory tract were reported for hamsters infected with the Omicron variant (Abdelnabi et al., 2022; McMahan et al., 2022). Therefore, an Omicron titration study in hamsters similar to what we describe here for the ancestral virus may be rewarding.

In summary, we determined the extremely high susceptibility of Golden Syrian hamsters to a SARS-CoV-2 infection, and defined the MID as 1*10^−3^ TCID_50_. A very close monitoring of pet Golden Syrian hamsters that are kept in households with COVID-19 patients is therefore strongly recommended, for instance by using rapid tests. COVID-19 patients should strictly avoid any direct or indirect contact to their pet hamsters. The optimal infection dose for drug efficacy studies was determined as 1*10^2^ TCID_50_. These conclusions not only apply to Golden Syrian hamsters, but likely also to Chinese hamsters (*Cricetulus griseus*) and Roborovski Dwarf hamsters (*Phodopus roborovskii*), since these two species have been shown to also be highly susceptible (Bertzbach et al., 2021; Trimpert et al., 2020).

## Materials and Methods

### Experimental Design

This study was performed to quantify the susceptibility of Golden Syrian hamsters to an infection with SARS-CoV-2. All SARS-CoV-2 experimental studies were performed in the biosafety level 3 facilities at the Friedrich-Loeffler-Institut, Insel Riems, Germany.

### Cell line and virus

Vero E6 cells (Cell Culture Collection in Veterinary Medicine, FLI) were cultured in minimal essential medium (MEM) containing 10% fetal calf serum (FCS) at 37°C and 5% CO2. SARS-CoV-2 isolate 2019_nCoV Muc-IMB-1 (accession number LR824570; (Wolfel et al., 2020)) was kindly provided by Bundeswehr Institute of Microbiology, Munich, Germany. Virus propagation was maintained in Vero E6 cells in DMEM supplemented with 2% FCS. Prior to inoculation into hamsters, the virus stock was sequenced by using a generic metagenomics sequencing workflow as described previously (Wylezich, Papa, Beer, & Hoper, 2018) with some modifications. For reverse-transcribing RNA into cDNA, SuperScriptIV First-Strand cDNA Synthesis System (Invitrogen, Germany) and the NEBNext Ultra II Non-Directional RNA Second Strand Synthesis Module (New England Biolabs, Germany) were used, and library quantification was done with the QIAseq Library Quant Assay Kit (Qiagen, Germany). Libraries were sequenced without applying further SARS-CoV-2 enrichment using an Ion 530 chip and chemistry for 400 base pair reads on an Ion Torrent S5XL instrument (Thermo Fisher Scientific, Germany).

### Virus titration - tissue culture infectious dose50

Samples were serially diluted in MEM containing 2% FCS and 100 Units Penicillin / 0.1 mg Streptomycin (P/S) (Millipore Sigma, Germany). Vero E6 cells were incubated with 100 µl of ten-fold dilutions of sample dilutions added in quadruplicates for 1 h at 37°C before 100 µl MEM containing 2% FCS and P/S were added per well and plates were incubated for 5 days at 37°C and 5% CO2. Supernatant was removed and cells were fixed with 4% formalin. Next, plates were stained with 1% crystal violet and titers were determined following the Spearman Kaerber method (Atkinson, 1961).

### Animal studies

Male Golden Syrian hamsters (*Mesocricetus auratus*), 5-7 weeks old with a body weight of 80 – 100 g, were obtained from Janvier Labs, France. Three hamsters were housed in individually ventilated cages (IVC). Animals had *ad libitum* access to food and water. The animals’ well-being and body weight were checked daily. Handling and sampling were performed starting with the uninfected group and continuing from the low dose groups to the high dose groups to minimize the contamination risk.

#### Inoculation routes

To determine the optimal inoculation route for SARS-CoV-2, we inoculated groups of eight hamsters under isoflurane anaesthesia in parallel by the intranasal and orotracheal route with 100 µl containing 1*10^5^ TCID_50_. For the orotracheal challenge, the inoculum was administered on the root of the tongue (Fig. 1a), ensuring aspiration of the inoculum with the following inhalation. Oral swab samples were collected in DMEM containing P/S daily, starting one day before inoculation. Nasal washes were collected under isoflurane anaesthesia at days 2, 4, 6, 9, 11 and 13 by flushing 200 µl PBS along the animal’s nose. At 14 dpi, hamsters were euthanized by deep isoflurane anaesthesia, cardiac exsanguination and cervical dislocation, and nasal conchae, trachea and lung samples were collected and stored at -80°C for virological analysis. Tissue samples were also stored in 4% neutral-buffered formalin for histopathological analysis. Serum was separated from the collected blood.

### Titration in hamsters

To define the MID, three sets of experiments were performed, as the endpoint in the first two studies was not reached. For animal welfare reasons, we continued the dilutions in the second and third experiment, including an overlap ensuring the comparability of result. First, hamsters were inoculated with a dose of 1*10^5^ TCID_50_ SARS-CoV-2 and with serial dilutions from 1*10^3^ to 1*10^−1^ TCID_50_. Oral swab samples were collected at 1, 3, 5, and 6 dpi, nasal washes were collected at 2 and 4 dpi as described above. At 7 dpi, hamsters were euthanized and sampled as described above.

In the second study, hamsters were inoculated with a serial dilution from 1*10^4^ to 1*10^−3^ TCID_50_ SARS-CoV-2. Sampling and autopsies were performed as described above.

In the third study, hamsters were inoculated with 1*10^2^ TCID_50_, and with serial dilutions from 1*10^0^ to 1*10^−4^ TCID_50_ SARS-CoV-2. This experiment was conducted for 10 days to allow the follow-up of clinical and virological data after a 2 - 3 day delayed disease onset in the groups inoculated with low doses. Oral swab samples were collected at 1, 3, 5, 6, 8 and 9 dpi, while nasal washes were collected at 2, 4 and 7 dpi. At 10 dpi, animals were sacrificed and autopsied as described above.

#### Ethical statement

Ethical approval for this study was obtained from the competent authority of the Federal State of Mecklenburg-Western Pomerania, Germany upon consultation with the Ethic Committee of Mecklenburg-Western Pomerania (file number: 7221.3-1.1-049/20), on the basis of national and European legislation, namely the EU council directive 2010/63/EU. Animal studies are continuously monitored by the Animal Welfare Officer and were approved by the Institutional Animal Care and Use Committee (IACUC).

### Total RNA extraction and SARS-CoV-2 detection

Total RNA was extracted from swab, nose fluid and tissue samples as described earlier (Schlottau et al., 2020). SARS-CoV-2 RNA was detected using “Envelope (E)-gene Sarbeco 6-carboxyfluorescein quantitative RT-PCR” (Corman et al., 2020) as described previously (Schlottau et al., 2020). Viral genome copy numbers were calculated from standard curves determined for 10^−2^ to 10^−5^ dilutions containing known copy numbers of SARS-CoV-2. Selected samples were analysed for the presence of subgenomic RNA (sgRNA) as an indication of virus replication, using a published protocol (Alexandersen et al., 2020; Hoffmann et al., 2021). Quantitative Realtime PCR was performed with the qScript XLT One-Step RT-qPCR ToughMix (QuantaBio/VWR). Primer sequences for the ORF 7a detection are available upon request.

### Indirect SARS-CoV-2 RBD ELISA

SARS-CoV-2 specific antibodies were detected using a published protocol (Wernike et al., 2021) with the modification of using a 1:30.000 dilution of Protein A/G (Thermo Fisher) in exchange of the multi-species conjugate.

To determine a cut-off-value and the diagnostic sensitivity of this modified assay, we tested 53 negative hamster sera and 227 sera of SARS-CoV-2 infected hamsters. The area under the receiver operating characteristic (ROC) curve was used to determine the ELISA cut-off-value. Statistical analyses were performed using MedCalc for Windows, version 19.4 (MedCalc Software, Ostend, Belgium). p-value < 0.01 was regarded as statistically significant.

### Lateral flow device (LFD) Rapid Test

Oral swab samples of groups inoculated with 1*10^3^ to 1*10^−2^ TCID_50_ SARS-CoV-2 were collected at 6 dpi in lysis buffer supplied with the Nowcheck COVID-19 LFD (concile GmbH, Freiburg, Germany) testkit. 120 µl of the suspension was applied on the LFD and incubated at RT for 15 min. We also analysed nasal wash samples collected at 7 dpi by diluting 25 µl of the fluid into 300 µl of the supplied lysis buffer and applying 120 μl of this mixture to the LFD. Evaluation of the control (C) and test (T) bands was performed according to the instructions. LFDs were imaged and densitometry was performed on the C and T bands using ImageJ (ImageJ 1.52a, Wayne Rasband, NIH, USA). These band quantifications were used to calculate a T/C ratio. Standard deviation was calculated from the T/C ratios within each respective group.

### Pathology of lung samples collected at 10 dpi

During autopsy, the percentage of dark red discoloration per total lung tissue was estimated. The left lung lobe was carefully removed, immersion-fixed in 10% neutral-buffered formalin, paraffin-embedded, and 2-3-μm sections were stained with hematoxylin and eosin (HE). Slides were scanned using a Hamamatsu S60 scanner, and evaluated using the NDPview.2 plus software (Version 2.8.24, Hamamatsu Photonics, K.K. Japan). The lung tissue was evaluated using a 500×500µm grid, and the extent of pneumonia-associated consolidation was recorded as percentage of affected lung fields. Further, the lung was examined for the presence of SARS-CoV2-characteristic lesions described for hamsters, i.e. intra-alveolar, interstitial, peribronchial and perivascular inflammatory infiltrates, alveolar edema, necrosis of the bronchial and alveolar epithelium, diffuse alveolar damage, vasculitis or endothelialitis, pneumocyte type 2 hyperplasia/hypertrophy with bronchialisation and atypical cells, and hypertrophy/hyperplasia of the bronchial epithelium. Archived lung tissues from hamsters infected with 1*10^4.5^ TCID_50_ were included as positive controls. Evaluation and interpretation were performed by a board-certified pathologist (DiplECVP) following a post examination masking approach (Meyerholz & Beck, 2018).

### Sequence analysis

Full genome sequences were generated via reference mapping with the Genome Sequencer software suite (version 2.6; Roche; default software settings for quality filtering and mapping), using SARS-CoV-2 strain 2019_nCoV_Muc_IMB1 (accession number LR824570) as reference. Consensus sequences and underlying sequence reads were visualized using Geneious Prime (10.2.3; Biomatters, Auckland, New Zealand). The presence of single nucleotide variants (SNVs) was checked using the variant analysis tool implemented in Geneious Prime (default settings, minimum variant frequency 0.02).

### Statistical analysis

Mean values determined for the experimental groups were compared using analysis of variance (ANOVA) with Tukey’s post-hoc tests for multiple comparisons and non-parametric Kruskal-Wallis test followed by the Dunn’s method for multiple comparisons. Data were analyzed using GraphPad Prism (version 9; GraphPad Software, Inc., CA, USA) and SPSS software (IBM Corp. Released 2011. IBM SPSS Statistics for Windows, Version 20.0, IBM Corporation, Armonk, NY, USA). *P*-value < 0.05 was considered statistically significant.

For histopathology, data were tested for Gaussian distribution using the Shapiro-Wilk test, followed by one-way ANOVA with Tukey’s post-hoc tests for multiple comparison.

## Acknowledgments

Daniel Balkema, Weda Hoffmann, Julia Neumeister, Silvia Schuparis, Emilie Stutz and Patrick Zitzow are thanked for their excellent technical assistance. Doreen Fiedler, Steffen Kiepert, Frank Klipp, Christian Lipinski and Harald Manthei are thanked for their support in the animal experiments.

## Funding

Max-Planck-Gesellschaft and Max-Planck-Förderstiftung FLI-internal funds The funders had no role in study design, data collection and analysis, decision to publish or preparation of the manuscript.

## Author contributions

Connceptualization: ABB, CB; Methodology: ABB, CB, MK; Formal Analysis: CB, BM, BS, AB; Investigation: ABB, CB, BM, AB, SW, CW; Visualization: CB, BM, AB; Supervision: ABB, DH; Writing—original draft: ABB, CB, BM; Writing—review & editing: TCM, MHG, DH, DG; Resources, DG, MHG, TCM

All authors have read and agreed to the published version of the manuscript.

## Competing interests

The authors declare they have no competing interests.

## Data and materials availability

All data needed to evaluate the conclusions in the paper are present in the paper and/or the Supplementary Materials.

---

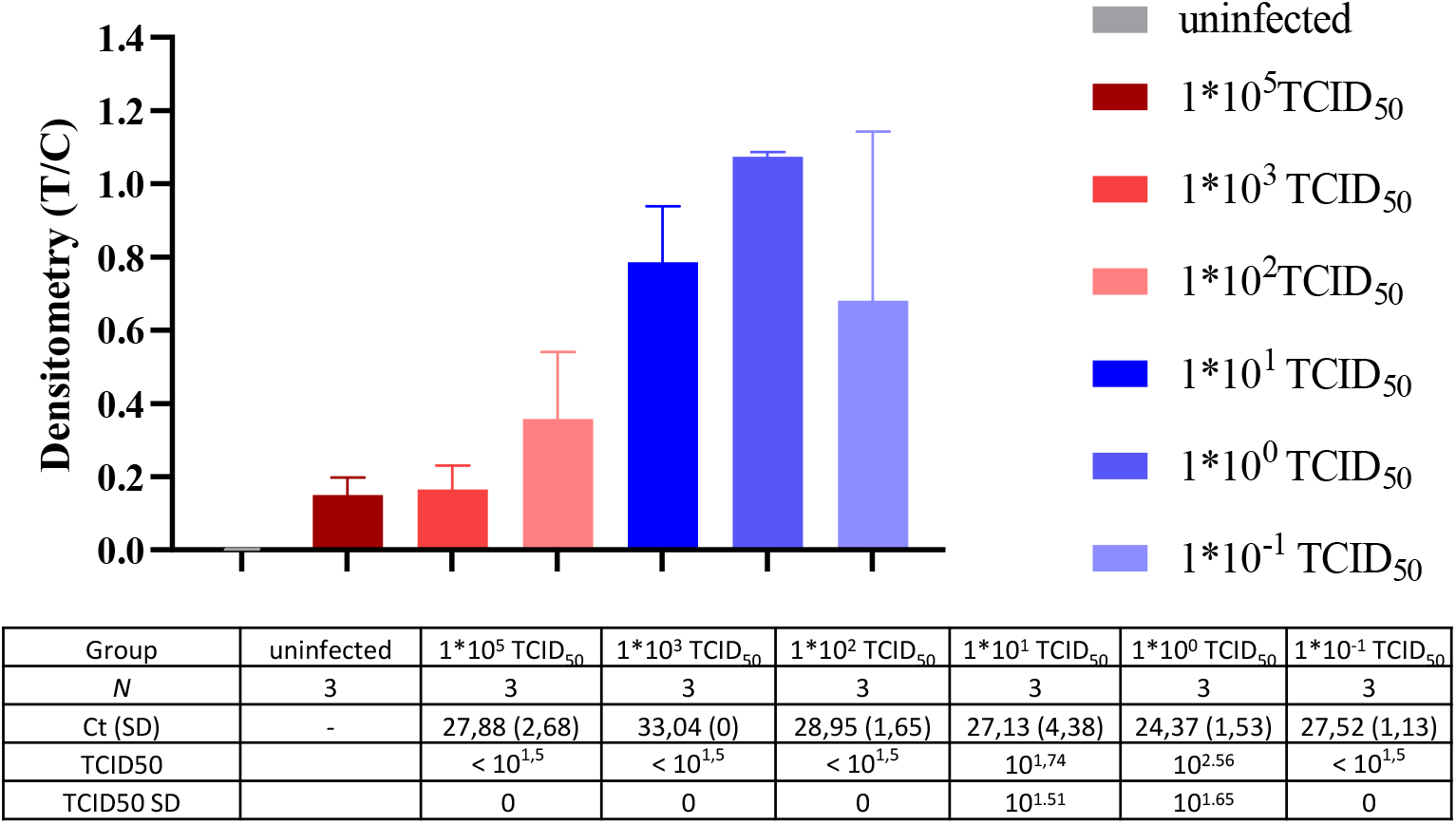

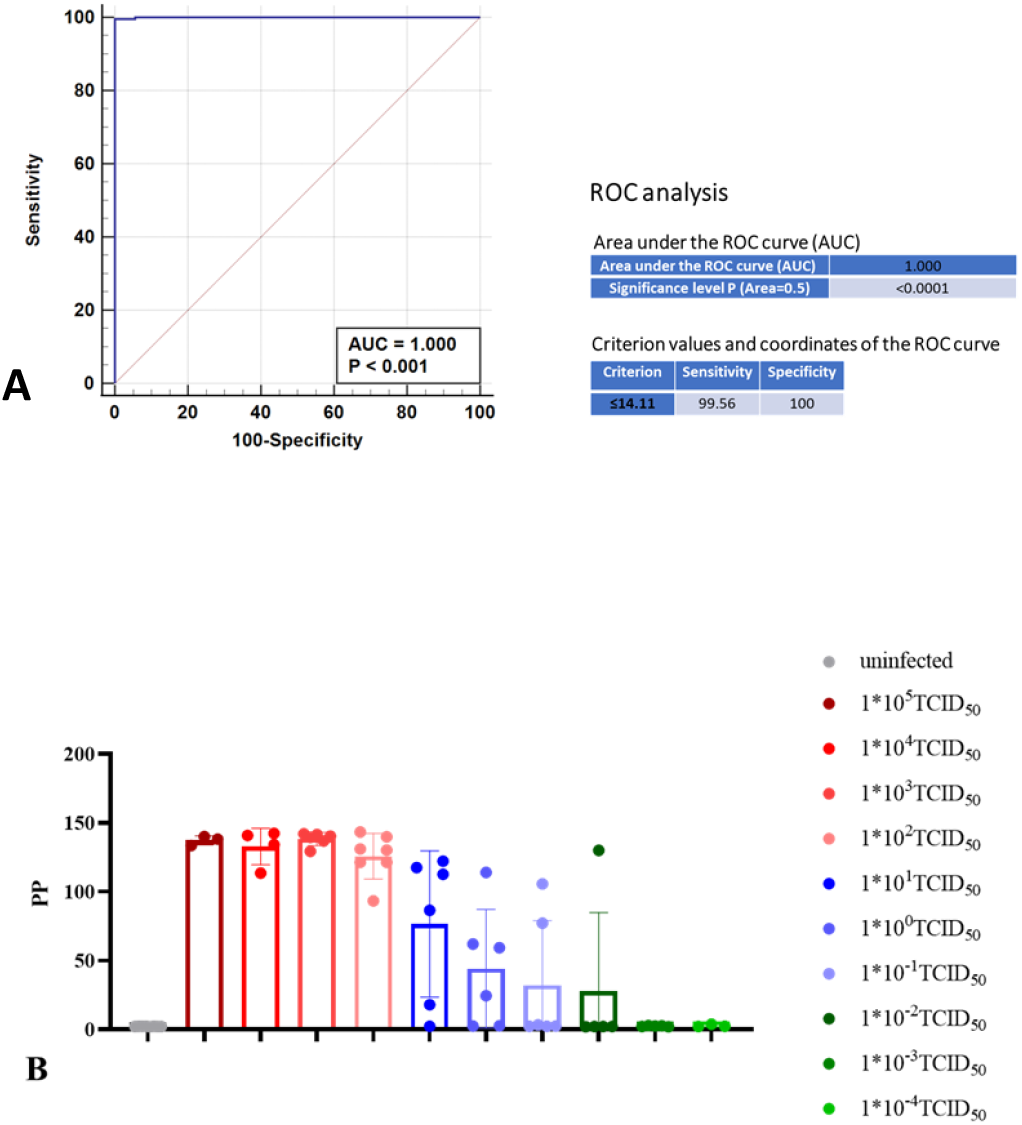

## Notes

### Competing Interest Statement

The authors have declared no competing interest.

## References

Abdelnabi, R., Foo, C. S., Zhang, X., Lemmens, V., Maes, P., Slechten, B., … Neyts, J. (2022). The omicron (B.1.1.529) SARS-CoV-2 variant of concern does not readily infect Syrian hamsters. Antiviral Res, 198, 105253. doi:10.1016/j.antiviral.2022.105253

Alexandersen, S., Chamings, A., & Bhatta, T. R. (2020). SARS-CoV-2 genomic and subgenomic RNAs in diagnostic samples are not an indicator of active replication. Nat Commun, 11(1), 6059. doi:10.1038/s41467-020-19883-7

Atkinson, G. F. (1961). The Spearman-Karber Method of Estimating <sub>50</sub>% Endpoints. Retrieved from https://hdl.handle.net/1813/32006

Basu, S. (2021). Computational characterization of inhaled droplet transport to the nasopharynx. Sci Rep, 11(1), 6652. doi:10.1038/s41598-021-85765-7

Bertzbach, L. D., Vladimirova, D., Dietert, K., Abdelgawad, A., Gruber, A. D., Osterrieder, N., & Trimpert, J. (2021). SARS-CoV-2 infection of Chinese hamsters (Cricetulus griseus) reproduces COVID-19 pneumonia in a well-established small animal model. Transboundary and Emerging Diseases, 68(3), 1075–1079. doi:https://doi.org/10.1111/tbed.13837

Chan, J. F., Zhang, A. J., Yuan, S., Poon, V. K., Chan, C. C., Lee, A. C., Yuen, K. Y. (2020). Simulation of the Clinical and Pathological Manifestations of Coronavirus Disease 2019 (COVID-19) in a Golden Syrian Hamster Model: Implications for Disease Pathogenesis and Transmissibility. Clin Infect Dis, 71(9), 2428–2446. doi:10.1093/cid/ciaa325

Cheng, C., Zhang, D., Dang, D., Geng, J., Zhu, P., Yuan, M., Duan, G. (2021). The incubation period of COVID-19: a global meta-analysis of 53 studies and a Chinese observation study of 11 545 patients. Infect Dis Poverty, 10(1), 119. doi:10.1186/s40249-021-00901-9

Chiba, S., Kiso, M., Nakajima, N., Iida, S., Maemura, T., Kuroda, M., Kawaoka, Y. (2022). Co-administration of Favipiravir and the Remdesivir Metabolite GS-441524 Effectively Reduces SARS-CoV-2 Replication in the Lungs of the Syrian Hamster Model. mBio, e0304421. doi:10.1128/mbio.03044-21

Corman, V. M., Landt, O., Kaiser, M., Molenkamp, R., Meijer, A., Chu, D. K., Drosten, C. (2020). Detection of 2019 novel coronavirus (2019-nCoV) by real-time RT-PCR. Euro Surveill, 25(3). doi:10.2807/1560-7917.ES.2020.25.3.2000045

Dowall, S., Salguero, F. J., Wiblin, N., Fotheringham, S., Hatch, G., Parks, S., … Hewson, R. (2021). Development of a Hamster Natural Transmission Model of SARS-CoV-2 Infection. Viruses, 13(11), 2251. Retrieved from https://www.mdpi.com/1999-4915/13/11/2251

Elias, C., Sekri, A., Leblanc, P., Cucherat, M., & Vanhems, P. (2021). The incubation period of COVID-19: A meta-analysis. Int J Infect Dis, 104, 708–710. doi:10.1016/j.ijid.2021.01.069

Francis, M. E., Goncin, U., Kroeker, A., Swan, C., Ralph, R., Lu, Y., Kelvin, A. A. (2021). SARS-CoV-2 infection in the Syrian hamster model causes inflammation as well as type I interferon dysregulation in both respiratory and non-respiratory tissues including the heart and kidney. PLoS Pathog, 17(7), e1009705. doi:10.1371/journal.ppat.1009705

Gruber, A. D., Firsching, T. C., Trimpert, J., & Dietert, K. (2021). Hamster models of COVID-19 pneumonia reviewed: How human can they be? Vet Pathol, 3009858211057197. doi:10.1177/03009858211057197

Haagmans, B. L., & Koopmans, M. P. G. (2022). Spreading of SARS-CoV-2 from hamsters to humans. Lancet, 399(10329), 1027–1028. doi:10.1016/S0140-6736(22)00423-8

Hoffmann, D., Corleis, B., Rauch, S., Roth, N., Muhe, J., Halwe, N. J., … Beer, M. (2021). CVnCoV and CV2CoV protect human ACE2 transgenic mice from ancestral B BavPat1 and emerging B.1.351 SARS-CoV-2. Nat Commun, 12(1), 4048. doi:10.1038/s41467-021-24339-7

Johnson, S., Martinez, C. I., Tedjakusuma, S. N., Peinovich, N., Dora, E. G., Birch, S. M., … Tucker, S. N. (2022). Oral Vaccination Protects Against Severe Acute Respiratory Syndrome Coronavirus 2 in a Syrian Hamster Challenge Model. J Infect Dis, 225(1), 34–41. doi:10.1093/infdis/jiab561

Karimzadeh, S., Bhopal, R., & Nguyen Tien, H. (2021). Review of infective dose, routes of transmission and outcome of COVID-19 caused by the SARS-COV-2: comparison with other respiratory viruses-CORRIGENDUM. Epidemiol Infect, 149, e116. doi:10.1017/S0950268821001084

Kok, K. H., Wong, S. C., Chan, W. M., Wen, L., Chu, A. W., Ip, J. D., … Yuen, K. Y. (2022). Co-circulation of two SARS-CoV-2 variant strains within imported pet hamsters in Hong Kong. Emerg Microbes Infect, 11(1), 689–698. doi:10.1080/22221751.2022.2040922

McMahan, K., Giffin, V., Tostanoski, L. H., Chung, B., Siamatu, M., Suthar, M. S., … Barouch, D. H. (2022). Reduced Pathogenicity of the SARS-CoV-2 Omicron Variant in Hamsters. bioRxiv, 2022.2001.2002.474743. doi:10.1101/2022.01.02.474743

Meseda, C. A., Stauft, C. B., Selvaraj, P., Lien, C. Z., Pedro, C., Nunez, I. A., … Weir, J. P. (2021). MVA vector expression of SARS-CoV-2 spike protein and protection of adult Syrian hamsters against SARS-CoV-2 challenge. NPJ Vaccines, 6(1), 145. doi:10.1038/s41541-021-00410-8

Meyerholz, D. K., & Beck, A. P. (2018). Principles and approaches for reproducible scoring of tissue stains in research. Lab Invest, 98(7), 844–855. doi:10.1038/s41374-018-0057-0

Mohandas, S., Yadav, P. D., Sapkal, G., Shete, A. M., Deshpande, G., Nyayanit, D. A., … Jain, R. (2022). Pathogenicity of SARS-CoV-2 Omicron in Syrian hamsters and its neutralization with different Variants of Concern. bioRxiv, 2022.2001.2019.477013. doi:10.1101/2022.01.19.477013

Mohandas, S., Yadav, P. D., Shete, A., Nyayanit, D., Sapkal, G., Lole, K., & Gupta, N. (2021). SARS-CoV-2 Delta Variant Pathogenesis and Host Response in Syrian Hamsters. Viruses, 13(9). doi:10.3390/v13091773

O’Donnell, K. L., Pinski, A. N., Clancy, C. S., Gourdine, T., Shifflett, K., Fletcher, P., … Marzi, A. (2021). Pathogenic and transcriptomic differences of emerging SARS-CoV-2 variants in the Syrian golden hamster model. EBioMedicine, 73, 103675. doi:10.1016/j.ebiom.2021.103675

Rosenke, K., Meade-White, K., Letko, M., Clancy, C., Hansen, F., Liu, Y., … Feldmann, H. (2020). Defining the Syrian hamster as a highly susceptible preclinical model for SARS-CoV-2 infection. bioRxiv. doi:10.1101/2020.09.25.314070

Schlottau, K., Rissmann, M., Graaf, A., Schon, J., Sehl, J., Wylezich, C., … Beer, M. (2020). SARS-CoV-2 in fruit bats, ferrets, pigs, and chickens: an experimental transmission study. Lancet Microbe, 1(5), e218–e225. doi:10.1016/S2666-5247(20)30089-6

Sia, S. F., Yan, L.-M., Chin, A. W. H., Fung, K., Choy, K.-T., Wong, A. Y. L., … Yen, H.-L. (2020). Pathogenesis and transmission of SARS-CoV-2 in golden hamsters. Nature, 583(7818), 834–838. doi:10.1038/s41586-020-2342-5

Taylor, R., Bowen, R., Demarest, J. F., DeSpirito, M., Hartwig, A., Bielefeldt-Ohmann, H., … Babu, Y. S. (2021). Activity of Galidesivir in a Hamster Model of SARS-CoV-2. Viruses, 14(1). doi:10.3390/v14010008

Trimpert, J., Vladimirova, D., Dietert, K., Abdelgawad, A., Kunec, D., Dökel, S., … Osterrieder, N. (2020). The Roborovski Dwarf Hamster Is A Highly Susceptible Model for a Rapid and Fatal Course of SARS-CoV-2 Infection. Cell Reports, 33(10), 108488. doi:https://doi.org/10.1016/j.celrep.2020.108488

Wernike, K., Aebischer, A., Michelitsch, A., Hoffmann, D., Freuling, C., Balkema-Buschmann, A., … Beer, M. (2021). Multi-species ELISA for the detection of antibodies against SARS-CoV-2 in animals. Transbound Emerg Dis, 68(4), 1779–1785. doi:10.1111/tbed.13926

Wolfel, R., Corman, V. M., Guggemos, W., Seilmaier, M., Zange, S., Muller, M. A., … Wendtner, C. (2020). Virological assessment of hospitalized patients with COVID-2019. Nature, 581(7809), 465–469. doi:10.1038/s41586-020-2196-x

Wylezich, C., Papa, A., Beer, M., & Hoper, D. (2018). A Versatile Sample Processing Workflow for Metagenomic Pathogen Detection. Sci Rep, 8(1), 13108. doi:10.1038/s41598-018-31496-1

Yadav, P. A.-O., Mendiratta, S. K., Mohandas, S., Singh, A. K., Abraham, P., Shete, A., … Jain, M. ZRC3308 Monoclonal Antibody Cocktail Shows Protective Efficacy in Syrian Hamsters against SARS-CoV-2 Infection. LID - 10.3390/v13122424 [doi] LID - 2424. (1999-4915 (Electronic)).

Yen, H.-L. a. S., Thomas HC and Brackman, Christopher J. and Chuk, Shirley SY and Cheng, Samuel M.S. and Gu, Haogao and Chang, Lydia DJ and Krishnan, Pavithra and Ng, Daisy YM and Liu, Gigi YZ and Hui, Mani MY and Ho, Sin Ying and Tam, Karina WS and Law, Pierra YT and Su, Wen and Sia, Sin Fun and Choy, Ka-Tim and Cheuk, Sammi SY and Lau, Sylvia PN and Tang, Amy WY and Koo, Joe CT and Yung, Louise and Leung, Gabriel and Peiris, J.S. Malik and Poon, Leo LM. (2022). Transmission of SARS-CoV-2 (Variant Delta) from Pet Hamsters to Humans and Onward Human Propagation of the Adapted Strain: A Case Study. The Lancet, [Preprint]. doi:http://dx.doi.org/10.2139/ssrn.4017393.

