## Supplementary material for "Compellingly high SARS-CoV-2 susceptibility of Golden Syrian hamsters suggests multiple zoonotic infections of pet hamsters during the COVID-19 pandemic": Figure S1

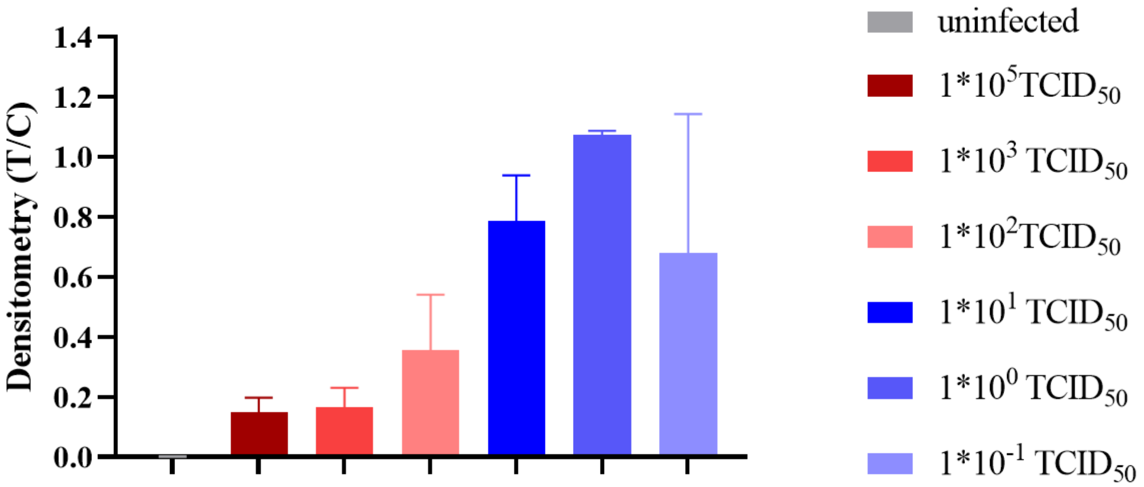


| Group | uninfected | 1*10⁵ $TCID_{50}$ | 1*10³ $TCID_{50}$ | 1*10² $TCID_{50}$ | 1*10¹ $TCID_{50}$ | 1*10⁰ $TCID_{50}$ | 1*10⁻¹ $TCID_{50}$ |
|---|---|---|---|---|---|---|---|
| *N* | 3 | 3 | 3 | 3 | 3 | 3 | 3 |
| Ct (SD) | - | 27,88 (2,68) | 33,04 (0) | 28,95 (1,65) | 27,13 (4,38) | 24,37 (1,53) | 27,52 (1,13) |
| TCID50 | | < $10^{1,5}$ | < $10^{1,5}$ | < $10^{1,5}$ | $10^{1,74}$ | $10^{2.56}$ | < $10^{1,5}$ |
| TCID50 SD | | 0 | 0 | 0 | $10^{1.51}$ | $10^{1.65}$ | 0 |

**Figure S 1**. **Analysis of oral swab samples by NowCheck COVID-19 Ag Test (LFD).** Qualitative results of antigen assay after analysis of animals infected with doses of $10^5$ to $10^{-1}$ $TCID_{50}$.
