## Supplementary material for "Compellingly high SARS-CoV-2 susceptibility of Golden Syrian hamsters suggests multiple zoonotic infections of pet hamsters during the COVID-19 pandemic": Figure S2

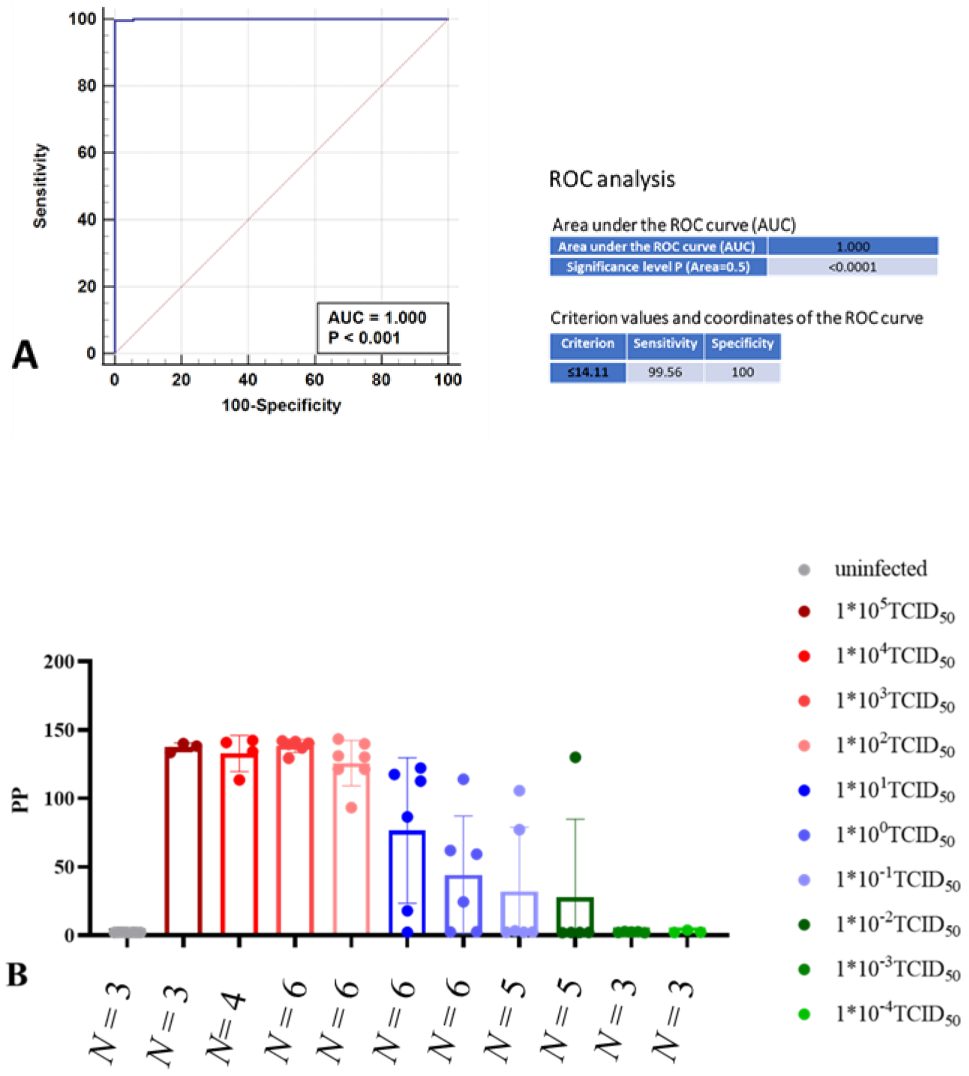


**Fig. S2. SARS-CoV-2 RBD-ELISA results of 280 hamster sera.** (A) ROC analysis using 53 negative hamster sera and 227 sera of experimentally SARS-CoV-2 infected hamsters. Diagnostic sensitivity of the SARS-CoV-2 RBD ELISA is 99.56% (95% CI 95.9–99.9) and diagnostic specificity is 100% (95% CI 87.7–100) with AUC 0.999 (p-value <0.01). (B) Dose-dependent development of SARS-CoV-2 specific antibodies within 7-10 days in groups infected with a dose of $1*10^{-2}$ $TCID_{50}$ or higher, with the variability within the groups increasing at infection doses below $1*10^{2}$ $TCID_{50,}$ as shown by the standard error of the mean (SEM).
