## Supplementary material for "Compellingly high SARS-CoV-2 susceptibility of Golden Syrian hamsters suggests multiple zoonotic infections of pet hamsters during the COVID-19 pandemic": Table S1

### Table S1. Significance levels calculated for the different infection dose groups.

| Body weight (%) | | | | | | | | | | Oral swab samples (Realtime PCR) | | |
|---|---|---|---|---|---|---|---|---|---|---|---|---|
| 1 dpi | 2 dpi | 3 dpi | 4 dpi | 5 dpi | 6 dpi | 7 dpi | 8 dpi | 9 dpi | 10 dpi | 6 dpi | 8 dpi | 9 dpi |
| <0.05* | <0.05* | <0.05* | <0.05* | <0.05* | <0.05* | <0.05* | <0.05* | <0.05* | <0.05* | <0.05* | <0.05* | <0.05* |
| Un vs. $10^5$ | Un vs. $10^4$ | Un vs. $10^4$ | Un vs. $10^5$ | Un vs. $10^5$ | Un vs. $10^5$ | Un vs. $10^5$ | Un vs. $10^2$ | Un vs. $10^2$ | Un vs. $10^2$ | $10^5$ vs. $10^{-2}$ | $10^2$ vs. $10^{-1}$ | $10^2$ vs. $10^{-1}$ |
| $10^5$ vs. $10^{-2}$ | Un vs. $10^{-3}$ | Un vs. $10^2$ | Un vs. $10^4$ | Un vs. $10^4$ | Un vs. $10^4$ | Un vs. $10^4$ | Un vs. 10 | Un vs. 10 | Un vs. 10 | $10^4$ vs. $10^{-2}$ | | |
| $10^4$ vs. $10^{-3}$ | $10^4$ vs. $10^1$ | $10^5$ vs. $10^{-1}$ | Un vs. $10^3$ | Un vs. $10^3$ | Un vs. $10^3$ | Un vs. $10^3$ | $10^2$ vs. $10^{-2}$ | $10^2$ vs. $10^{-1}$ | $10^2$ vs. $10^{-1}$ | $10^3$ vs. $10^{-2}$ | | |
| $10^3$ vs. $10^{-2}$ | $10^4$ vs. 10 | $10^4$ vs. $10^1$ | Un vs. $10^2$ | Un vs. $10^2$ | Un vs. $10^2$ | Un vs. $10^2$ | $10^2$ vs. $10^{-3}$ | $10^2$ vs. $10^{-2}$ | $10^2$ vs. $10^{-2}$ | $10^2$ vs. $10^{-2}$ | | |
| $10^2$ vs. $10^{-3}$ | $10^4$ vs. $10^{-1}$ | $10^4$ vs. 10 | $10^5$ vs. $10^{-1}$ | $10^5$ vs. $10^{-1}$ | $10^5$ vs. $10^{-1}$ | $10^5$ vs. $10^{-1}$ | $10^2$ vs. $10^{-4}$ | $10^2$ vs. $10^{-3}$ | $10^2$ vs. $10^{-3}$ | $10^1$ vs. $10^{-2}$ | | |
| $10^1$ vs. $10^{-3}$ | $10^3$ vs. $10^{-3}$ | $10^4$ vs. $10^{-1}$ | $10^4$ vs. $10^1$ | $10^5$ vs. $10^{-2}$ | $10^5$ vs. $10^{-2}$ | $10^5$ vs. $10^{-2}$ | 10 vs. $10^{-2}$ | $10^2$ vs. $10^{-4}$ | $10^2$ vs. $10^{-4}$ | 10 vs. $10^{-2}$ | | |
| 10 vs. $10^{-3}$ | $10^2$ vs. $10^{-1}$ | $10^4/10^{-2}$ | $10^4$ vs. 10 | $10^4$ vs. $10^1$ | $10^5$ vs. $10^{-3}$ | $10^4$ vs. $10^{-1}$ | 10 vs. $10^{-3}$ | 10 vs. $10^{-2}$ | 10 vs. $10^{-2}$ | $10^{-1}$ vs. $10^{-2}$ | | |
| $10^{-1}$ vs. $10^{-3}$ | $10^1$ vs. $10^4$ | $10^4/10^{-3}$ | $10^4$ vs. $10^{-1}$ | $10^4$ vs. 10 | $10^5$ vs. $10^{-4}$ | $10^4$ vs. $10^{-2}$ | 10 vs. $10^{-4}$ | 10 vs. $10^{-3}$ | 10 vs. $10^{-3}$ | $10^{-3}$ vs. $10^{-2}$ | | |
| | $10^1$ vs. $10^{-3}$ | $10^4/10^{-4}$ | $10^4$ vs. $10^{-2}$ | $10^4$ vs. $10^{-1}$ | $10^4$ vs. $10^1$ | $10^4$ vs. $10^{-3}$ | $10^{-1}$ vs. $10^{-2}$ | 10 vs. $10^{-4}$ | 10 vs. $10^{-4}$ | | | |
| | 10 vs. $10^{-3}$ | $10^3/10^{-1}$ | $10^4$ vs. $10^{-3}$ | $10^4$ vs. $10^{-2}$ | $10^4$ vs. 10 | $10^4$ vs. $10^{-4}$ | | $10^{-1}$ vs. $10^{-2}$ | $10^{-1}$ vs. $10^{-2}$ | | | |
| | $10^{-1}$ vs. $10^{-3}$ | $10^2/10^1$ | $10^4$ vs. $10^{-4}$ | $10^4$ vs. $10^{-3}$ | $10^4$ vs. $10^{-1}$ | $10^2$ vs. $10^{-1}$ | | $10^{-1}$ vs. $10^{-3}$ | $10^{-1}$ vs. $10^{-3}$ | | | |
| | $10^{-2}$ vs. $10^{-3}$ | $10^2/10$ | $10^3$ vs. $10^{-1}$ | $10^4$ vs. $10^{-4}$ | $10^4$ vs. $10^{-2}$ | $10^2$ vs. $10^{-2}$ | | $10^{-1}$ vs. $10^{-4}$ | $10^{-1}$ vs. $10^{-4}$ | | | |
| | | $10^2/10^{-1}$ | $10^2$ vs. 10 | $10^3$ vs. $10^{-1}$ | $10^4$ vs. $10^{-3}$ | | | | | | | |
| | | $10^2/10^{-2}$ | $10^2$ vs. $10^{-1}$ | $10^2$ vs. $10^{-1}$ | $10^4$ vs. $10^{-4}$ | | | | | | | |
| | | | $10^2$ vs. $10^{-2}$ | $10^2$ vs. $10^{-2}$ | $10^3$ vs. $10^{-1}$ | | | | | | | |
| | | | $10^2$ vs. $10^{-3}$ | $10^2$ vs. $10^{-3}$ | $10^2$ vs. $10^{-1}$ | | | | | | | |
| | | | | | $10^2$ vs. $10^{-2}$ | | | | | | | |
| | | | | | $10^2$ vs. $10^{-3}$ | | | | | | | |

| Nasal washings (Realtime PCR) | | | Organs (7 dpi – Realtime PCR) | | |
|---|---|---|---|---|---|
| 2 dpi | 4 dpi | 7 dpi | Nasal conchae | Trachea | Lung |
| <0.05* | <0.05* | <0.05* | <0.05* | <0.05* | <0.05* |
| $10^5$ vs. 10 | $10^5$ vs. $10^{-2}$ | $10^2$ vs. $10^{-2}$ | $10^3$ vs. $10^{-2}$ | $10^4$ vs. $10^{-3}$ | $10^5$ vs. $10^{-2}$ |
| $10^5$ vs. $10^{-1}$ | $10^5$ vs. $10^{-3}$ | $10^2$ vs. $10^{-3}$ | $10^1$ vs. $10^{-2}$ | $10^3$ vs. $10^{-3}$ | $10^4$ vs. $10^{-2}$ |
| $10^5$ vs. $10^{-2}$ | $10^5$ vs. $10^{-4}$ | 10 vs. $10^{-2}$ | 10 vs. $10^{-2}$ | | $10^3$ vs. $10^0$ |
| $10^5$ vs. $10^{-3}$ | $10^4$ vs. $10^{-2}$ | 10 vs. $10^{-3}$ | $10^{-1}$ vs. $10^{-2}$ | | $10^3$ vs. $10^{-1}$ |
| $10^5$ vs. $10^{-4}$ | $10^4$ vs. $10^{-3}$ | $10^{-1}$ vs. $10^{-2}$ | | | $10^3$ vs. $10^{-2}$ |
| $10^4$ vs. 10 | $10^4$ vs. $10^{-4}$ | $10^{-2}$ vs. $10^{-3}$ | | | $10^3$ vs. $10^{-3}$ |
| $10^4$ vs. $10^{-1}$ | $10^3$ vs. $10^{-2}$ | | | | $10^2$ vs. $10^0$ |
| $10^4$ vs. $10^{-2}$ | $10^3$ vs. $10^{-3}$ | | | | $10^2$ vs. $10^{-2}$ |
| $10^4$ vs. $10^{-3}$ | $10^3$ vs. $10^{-4}$ | | | | $10^1$ vs. $10^{-2}$ |
| $10^4$ vs. $10^{-4}$ | $10^2$ vs. $10^{-2}$ | | | | $10^0$ vs. $10^{-2}$ |
| $10^3$ vs. 10 | $10^2$ vs. $10^{-3}$ | | | | $10^{-1}$ vs. $10^{-2}$ |
| $10^3$ vs. $10^{-1}$ | $10^2$ vs. $10^{-4}$ | | | | $10^{-2}$ vs. $10^{-3}$ |
| $10^3$ vs. $10^{-2}$ | $10^1$ vs. $10^{-2}$ | | | | |
| $10^3$ vs. $10^{-3}$ | $10^1$ vs. $10^{-3}$ | | | | |
| $10^3$ vs. $10^{-4}$ | $10^1$ vs. $10^{-4}$ | | | | |
| $10^2$ vs. 10 | 10 vs. $10^{-2}$ | | | | |
| $10^2$ vs. $10^{-1}$ | 10 vs. $10^{-3}$ | | | | |
| $10^2$ vs. $10^{-2}$ | 10 vs. $10^{-4}$ | | | | |
| $10^2$ vs. $10^{-3}$ | $10^{-1}$ vs. $10^{-2}$ | | | | |
| $10^2$ vs. $10^{-4}$ | $10^{-2}$ vs. $10^{-3}$ | | | | |
| | $10^{-3}$ vs. $10^{-4}$ | | | | |
