## Supplementary material for "Compellingly high SARS-CoV-2 susceptibility of Golden Syrian hamsters suggests multiple zoonotic infections of pet hamsters during the COVID-19 pandemic": Table S2

**Table S2. PCR detection of SARS-CoV-2 sgRNA. N-gene RNA, standardized quantity (SQ) of N gene copy numbers. as well as $TCID_{50}$ for oral swab, nasal wash and tissue samples. Colors match and represent results from the same hamsters as shown in figures.**

| Infection dose | Sample | dpi | ct sg RNA | Ct N-gene | SQ N-gene | $TCID_{50}$ |
|---|---|---|---|---|---|---|
| 1 x $10^1$ $TCID_{50}$ | nasal wash | 2 | 32.88 | 29.14 | 1.60E+03 | <10^1.5 |
| 1 x $10^1$ $TCID_{50}$ | nasal wash | 2 | 22.52 | 21.60 | 2.83E+05 | <10^1.5 |
| 1 x $10^1$ $TCID_{50}$ | nasal wash | 2 | 25.18 | 22.90 | 1.16E+05 | 10^5.25 |
| 1 x $10^0$ $TCID_{50}$ | nasal wash | 2 | 32.87 | 29.16 | 1.57E+03 | <10^1.5 |
| 1 x $10^0$ $TCID_{50}$ | nasal wash | 2 | 23.19 | 20.26 | 7.11E+05 | 10^5 |
| 1 x $10^0$ $TCID_{50}$ | nasal wash | 2 | 28.53 | 23.50 | 7.67E+04 | 10^3 |
| 1 x $10^{-1}$ $TCID_{50}$ | nasal wash | 2 | 27.27 | 24.42 | 4.07E+04 | 10^2.25 |
| 1 x $10^{-1}$ $TCID_{50}$ | nasal wash | 2 | 39.87 | 41.14 | 4.23E-01 | <10^1.5 |
| 1 x $10^{-1}$ $TCID_{50}$ | nasal wash | 2 | 27.89 | 22.69 | 1.34E+05 | 10^2.5 |
| 1 x $10^{-2}$ $TCID_{50}$ | nasal wash | 2 | neg | neg | N/A | - |
| 1 x $10^{-2}$ $TCID_{50}$ | nasal wash | 2 | neg | neg | N/A | - |
| 1 x $10^{-3}$ $TCID_{50}$ | nasal wash | 2 | 18.59 | neg | N/A | - |
| 1 x $10^{-3}$ $TCID_{50}$ | nasal wash | 2 | neg | neg | N/A | - |
| 1 x $10^{-3}$ $TCID_{50}$ | nasal wash | 2 | neg | neg | N/A | - |
| 1 x $10^1$ $TCID_{50}$ | nasal wash | 4 | 30.50 | 23.77 | 1.51E+04 | <10^1.5 |
| 1 x $10^1$ $TCID_{50}$ | nasal wash | 4 | 21.44 | 16.34 | 1.89E+06 | 10^5.5 |
| 1 x $10^1$ $TCID_{50}$ | nasal wash | 4 | 25. Jan | 20.16 | 1.57E+05 | 10^3.5 |
| 1 x $10^0$ $TCID_{50}$ | nasal wash | 4 | 17. Apr | 14.33 | 6.97E+06 | 10^6.75 |
| 1 x $10^0$ $TCID_{50}$ | nasal wash | 4 | 16.41 | Dez 69 | 2.03E+07 | 10^6.25 |
| 1 x $10^0$ $TCID_{50}$ | nasal wash | 4 | 17. Sep | Dez 23 | 2.73E+07 | 10^6.5 |
| 1 x $10^{-1}$ $TCID_{50}$ | nasal wash | 4 | 18.78 | 16.16 | 2.12E+06 | 10^5.5 |
| 1 x $10^{-1}$ $TCID_{50}$ | nasal wash | 4 | 23.43 | 18.26 | 5.41E+05 | 10^5 |
| 1 x $10^{-1}$ $TCID_{50}$ | nasal wash | 4 | 23.78 | 18.57 | 4.43E+05 | 10^4.25 |
| 1 x $10^{-2}$ $TCID_{50}$ | nasal wash | 4 | N/A | 36.98 | 2.82E+00 | <10^1.5 |
| 1 x $10^{-2}$ $TCID_{50}$ | nasal wash | 4 | 37.29 | 37.41 | 2.13E+00 | 10^6 |
| 1 x $10^{-3}$ $TCID_{50}$ | nasal wash | 4 | 24.45 | 24. Apr | 1.26E+04 | 10^2.75 |
| 1 x $10^{-3}$ $TCID_{50}$ | nasal wash | 4 | 34.05 | 29.29 | 4.18E+02 | <10^1.5 |
| 1 x $10^{-3}$ $TCID_{50}$ | nasal wash | 4 | N/A | 37.61 | 1.45E+00 | <10^1.5 |
| 1 x $10^{-2}$ $TCID_{50}$ | oral swab | 6 | . | 35.19 | 9.34E+00 | <10^1.5 |
| 1 x $10^{-2}$ $TCID_{50}$ | oral swab | 6 | N/A | 36.89 | 2.98E+00 | <10^1.5 |
| 1 x $10^{-3}$ $TCID_{50}$ | oral swab | 6 | 28.56 | 23.77 | 1.99E+04 | 10^2 |
| 1 x $10^{-3}$ $TCID_{50}$ | oral swab | 6 | 29.24 | 24.70 | 1.07E+04 | <10^1.5 |
| 1 x $10^{-3}$ $TCID_{50}$ | oral swab | 6 | 32.73 | 28.75 | 7.03E+02 | <10^1.5 |
| 1 x $10^2$ $TCID_{50}$ | nasal conchae | 7 | 19.67 | 14.57 | 1.06E+07 | 10^3.5 |
| 1 x $10^2$ $TCID_{50}$ | trachea | 7 | 34.16 | 29.98 | 2.56E+02 | <10^1.5 |
| 1 x $10^2$ $TCID_{50}$ | lung | 7 | 21.71 | 19.98 | 2.67E+05 | 10^2 |
| 1 x $10^2$ $TCID_{50}$ | nasal conchae | 7 | 22.64 | 18.17 | 9.14E+05 | 10^1.75 |

| | | | | | | |
|---|---|---|---|---|---|---|
| 1 x $10^{2}$ $TCID_{50}$ | trachea | 7 | 35.18 | 31.72 | 8.93E+01 | <10^1.5 |
| 1 x $10^{2}$ $TCID_{50}$ | lung | 7 | 25.95 | 23.74 | 2.06E+04 | <10^1.5 |
| 1 x $10^{2}$ $TCID_{50}$ | nasal conchae | 7 | 23.93 | 19.86 | 2.89E+05 | <10^1.5 |
| 1 x $10^{2}$ $TCID_{50}$ | trachea | 7 | 36.89 | 33.16 | 3.03E+01 | <10^1.5 |
| 1 x $10^{2}$ $TCID_{50}$ | lung | 7 | 27.55 | 26.65 | 2.82E+03 | <10^1.5 |
| 1 x $10^{1}$ $TCID_{50}$ | nasal chonchae | 7 | 22.59 | 17.19 | 1.79E+06 | 10^3 |
| 1 x $10^{1}$ $TCID_{50}$ | trachea | 7 | 26.23 | 19.31 | 4.20E+05 | <10^1.5 |
| 1 x $10^{1}$ $TCID_{50}$ | lung | 7 | 21.77 | 20.34 | 2.09E+05 | <10^1.5 |
| 1 x $10^{1}$ $TCID_{50}$ | nasal conchae | 7 | 24.62 | 19.71 | 3.21E+05 | <10^1.5 |
| 1 x $10^{1}$ $TCID_{50}$ | trachea | 7 | 29.13 | 25.34 | 6.93E+03 | <10^1.5 |
| 1 x $10^{1}$ $TCID_{50}$ | lung | 7 | 29.50 | 27.15 | 2.02E+03 | <10^1.5 |
| 1 x $10^{1}$ $TCID_{50}$ | nasal conchae | 7 | 17.99 | 15.44 | 5.89E+06 | 10^8.5 |
| 1 x $10^{1}$ $TCID_{50}$ | trachea | 7 | 30.29 | 25.87 | 4.81E+03 | <10^1.5 |
| 1 x $10^{1}$ $TCID_{50}$ | lung | 7 | 21.54 | 20. Jan | 2.62E+05 | 10^2 |
| 1 x $10^{0}$ $TCID_{50}$ | nasal conchae | 7 | 19.89 | 17.28 | 1.68E+06 | 10^7.5 |
| 1 x $10^{0}$ $TCID_{50}$ | trachea | 7 | 31.63 | 39.35 | 4.91E-01 | <10^1.5 |
| 1 x $10^{0}$ $TCID_{50}$ | lung | 7 | 18.74 | 16.18 | 3.55E+06 | 10^4.5 |
| 1 x $10^{0}$ $TCID_{50}$ | nasal conchae | 7 | 25.91 | 22.14 | 6.13E+04 | 10^3.25 |
| 1 x $10^{0}$ $TCID_{50}$ | trachea | 7 | 30.47 | N/A | neg | neg |
| 1 x $10^{0}$ $TCID_{50}$ | lung | 7 | 19.47 | 15.70 | 2.47E+06 | 10^3.75 |
| 1 x $10^{0}$ $TCID_{50}$ | nasal conchae | 7 | 20.50 | 15. Jan | 3.93E+06 | 10^4.25 |
| 1 x $10^{0}$ $TCID_{50}$ | trachea | 7 | 33.35 | 30.46 | 1.86E+02 | 10^2.5 |
| 1 x $10^{0}$ $TCID_{50}$ | lung | 7 | 19.73 | 17.13 | 9.36E+05 | 10^3.5 |
| 1 x $10^{-1}$ $TCID_{50}$ | nasal conchae | 7 | 21.59 | 15.89 | 2.17E+06 | - |
| 1 x $10^{-1}$ $TCID_{50}$ | trachea | 7 | 39.46 | 36.75 | 1.58E+00 | <10^1.5 |
| 1 x $10^{-1}$ $TCID_{50}$ | lung | 7 | 22.31 | 18.70 | 3.24E+05 | 10^2.25 |
| 1 x $10^{-1}$ $TCID_{50}$ | nasal conchae | 7 | 22. Mai | 17.35 | 8.07E+05 | 10^2.5 |
| 1 x $10^{-1}$ $TCID_{50}$ | trachea | 7 | 25.20 | 19.50 | 1.88E+05 | <10^1.5 |
| 1 x $10^{-1}$ $TCID_{50}$ | lung | 7 | 18.42 | 15. Jun | 3.81E+06 | 10^7.25 |
| 1 x $10^{-1}$ $TCID_{50}$ | nasal conchae | 7 | 29.21 | 23.35 | 1.38E+04 | <10^1.5 |
| 1 x $10^{-1}$ $TCID_{50}$ | trachea | 7 | 32.33 | 29.88 | 2.75E+02 | 10^1.75 |
| 1 x $10^{-1}$ $TCID_{50}$ | lung | 7 | 21.59 | 18.80 | 3.03E+05 | 10^4.75 |
| 1 x $10^{-2}$ $TCID_{50}$ | nasal conchae | 7 | 22.77 | 17.54 | 7.09E+05 | 10^6.5 |
| 1 x $10^{-2}$ $TCID_{50}$ | trachea | 7 | 27.48 | 22.47 | 2.52E+04 | 10^3 |
| 1 x $10^{-2}$ $TCID_{50}$ | lung | 7 | 40.80 | 36.89 | 1.44E+00 | <10^1.5 |
| 1 x $10^{-2}$ $TCID_{50}$ | nasal conchae | 7 | 41.48 | 36.45 | 1.94E+00 | <10^1.5 |
| 1 x $10^{-2}$ $TCID_{50}$ | trachea | 7 | N/A | N/A | N/A | N/A |
| 1 x $10^{-2}$ $TCID_{50}$ | lung | 7 | N/A | 36.39 | 2.02E+00 | <10^1.5 |
| 1 x $10^{-3}$ $TCID_{50}$ | nasal conchae | 7 | 20. Dez | 15.27 | 3.30E+06 | 10^5 |
| 1 x $10^{-3}$ $TCID_{50}$ | trachea | 7 | 24.14 | 18.57 | 3.54E+05 | 10^3 |
| 1 x $10^{-3}$ $TCID_{50}$ | lung | 7 | 21. Nov | 17.52 | 7.19E+05 | <10^1.5 |
| 1 x $10^{-3}$ $TCID_{50}$ | nasal conchae | 7 | 26. Okt | 19.75 | 1.59E+05 | 10^3 |

| | | | | | | |
|---|---|---|---|---|---|---|
| 1 x $10^{-3}$ $TCID_{50}$ | trachea | 7 | 26.43 | 19.98 | 1.36E+05 | 10^2.75 |
| 1 x $10^{-3}$ $TCID_{50}$ | lunge | 7 | 19.13 | 15.14 | 3.61E+06 | <10^1.5 |
| 1 x $10^{-3}$ $TCID_{50}$ | nasal conchae | 7 | 22. Sep | 16. Aug | 1.91E+06 | 10^6 |
| 1 x $10^{-3}$ $TCID_{50}$ | trachea | 7 | 25.74 | 19.43 | 1.98E+05 | 10^3.75 |
| 1 x $10^{-3}$ $TCID_{50}$ | lung | 7 | 21.31 | 17.52 | 7.21E+05 | <10^1.5 |
