## Supplementary material for "Compellingly high SARS-CoV-2 susceptibility of Golden Syrian hamsters suggests multiple zoonotic infections of pet hamsters during the COVID-19 pandemic": Table S3

**Table S3. Significance levels calculated for RNA levels detected in the organ samples of different infection dose groups (supplementary information to Fig. 8).**

**ANOVA (n=42)**

Lung

| | Sum of Squares | df | Mean Square | F | Sig. |
|---|---|---|---|---|---|
| Between Groups | 69.192 | 8 | 8.649 | 11.629 | .000 |
| Within Groups | 24.543 | 33 | .744 | | |
| Total | 93.736 | 41 | | | |

This initial analysis reveals statistically significant differences between the groups as a whole. The table below displays the results of Multiple Comparisons between the individual groups (Tukey post hoc test).

**Multiple Comparisons**

For better reading, the infection doses were marked as follows: B: 1 x $10^5$ $TCID_{50}$, C: 1 x $10^3$ $TCID_{50}$, D: 1 x $10^2$ $TCID_{50}$, E: 1 x $10^1$ $TCID_{50}$, F: 1 x $10^0$ $TCID_{50}$, G: 1 x $10^{-1}$ $TCID_{50}$, H: 1 x $10^{-2}$ $TCID_{50}$, I: 1 x $10^{-3}$ $TCID_{50}$, J: 1 x $10^{-4}$ $TCID_{50}$. *P*-values below 0.05 were considered statistically significant and are marked in green.

| (I) Dose | (J) Dose | Mean Difference (I-J) | Std. Error | *P* | 95% Confidence Interval | |
|---|---|---|---|---|---|---|
| | | | | | Lower Bound | Upper Bound |
| B | C | -1.35965 | .65867 | .513 | -3.5434 | .8241 |
| | D | .02835 | .60981 | 1.000 | -1.9934 | 2.0501 |
| | E | -.23524 | .60981 | 1.000 | -2.2570 | 1.7865 |
| | F | -.96856 | .60981 | .804 | -2.9903 | 1.0532 |
| | G | -1.96976 | .60981 | .061 | -3.9915 | .0520 |
| | H | -1.77085 | .60981 | .124 | -3.7926 | .2509 |
| | I | 3.69167* | .78726 | .001 | 1.0816 | 6.3017 |
| | J | -2.16720 | .70415 | .086 | -4.5017 | .1673 |
| C | B | 1.35965 | .65867 | .513 | -.8241 | 3.5434 |
| | D | 1.38801 | .55668 | .271 | -.4576 | 3.2336 |
| | E | 1.12441 | .55668 | .541 | -.7212 | 2.9700 |
| | F | .39110 | .55668 | .998 | -1.4545 | 2.2367 |
| | G | -.61011 | .55668 | .971 | -2.4557 | 1.2355 |
| | H | -.41120 | .55668 | .998 | -2.2568 | 1.4344 |
| | I | 5.05132* | .74686 | .000 | 2.5752 | 7.5275 |
| | J | -.80754 | .65867 | .945 | -2.9913 | 1.3762 |

| | | | | | | |
|---|---|---|---|---|---|---|
| D | B | -.02835 | .60981 | 1.000 | -2.0501 | 1.9934 |
| | C | -1.38801 | .55668 | .271 | -3.2336 | .4576 |
| | E | -.26359 | .49791 | 1.000 | -1.9144 | 1.3872 |
| | F | -.99691 | .49791 | .553 | -2.6477 | .6539 |
| | G | -1.99811* | .49791 | .009 | -3.6489 | -.3474 |
| | H | -1.79920* | .49791 | .024 | -3.4500 | -.1484 |
| | I | 3.66332* | .70415 | .000 | 1.3288 | 5.9978 |
| | J | -2.19555* | .60981 | .025 | -4.2173 | -.1738 |
| E | B | .23524 | .60981 | 1.000 | -1.7865 | 2.2570 |
| | C | -1.12441 | .55668 | .541 | -2.9700 | .7212 |
| | D | .26359 | .49791 | 1.000 | -1.3872 | 1.9144 |
| | F | -.73331 | .49791 | .860 | -2.3841 | .9174 |
| | G | -1.73452* | .49791 | .033 | -3.3853 | -.0838 |
| | H | -1.53561 | .49791 | .085 | -3.1864 | .1151 |
| | I | 3.92691* | .70415 | .000 | 1.5924 | 6.2614 |
| | J | -1.93196 | .60981 | .070 | -3.9537 | .0898 |
| F | B | .96856 | .60981 | .804 | -1.0532 | 2.9903 |
| | C | -.39110 | .55668 | .998 | -2.2367 | 1.4545 |
| | D | .99691 | .49791 | .553 | -.6539 | 2.6477 |
| | E | .73331 | .49791 | .860 | -.9174 | 2.3841 |
| | G | -1.00121 | .49791 | .547 | -2.6520 | .6496 |
| | H | -.80230 | .49791 | .792 | -2.4531 | .8485 |
| | I | 4.66022* | .70415 | .000 | 2.3257 | 6.9948 |
| | J | -1.19864 | .60981 | .576 | -3.2204 | .8231 |
| G | B | 1.96976 | .60981 | .061 | -.0520 | 3.9915 |
| | C | .61011 | .55668 | .971 | -1.2355 | 2.4557 |
| | D | 1.99811* | .49791 | .009 | .3474 | 3.6489 |
| | E | 1.73452* | .49791 | .033 | .0838 | 3.3853 |
| | F | 1.00121 | .49791 | .547 | -.6496 | 2.6520 |
| | H | .19891 | .49791 | 1.000 | -1.4518 | 1.8497 |
| | I | 5.66143* | .70415 | .000 | 3.3269 | 7.9960 |
| | J | -.19744 | .60981 | 1.000 | -2.2192 | 1.8243 |
| H | B | 1.77085 | .60981 | .124 | -.2509 | 3.7926 |
| | C | .41120 | .55668 | .998 | -1.4344 | 2.2568 |
| | D | 1.79920* | .49791 | .024 | .1484 | 3.4500 |
| | E | 1.53561 | .49791 | .085 | -.1151 | 3.1864 |
| | F | .80230 | .49791 | .792 | -.8485 | 2.4531 |
| | G | -.19891 | .49791 | 1.000 | -1.8497 | 1.4518 |
| | I | 5.46252* | .70415 | .000 | 3.1280 | 7.7970 |
| | J | -.39635 | .60981 | .999 | -2.4181 | 1.6254 |

| | | | | | | |
|---|---|---|---|---|---|---|
| I | B | -3.69167* | .78726 | .001 | -6.3017 | -1.0816 |
| | C | -5.05132* | .74686 | .000 | -7.5275 | -2.5752 |
| | D | -3.66332* | .70415 | .000 | -5.9978 | -1.3288 |
| | E | -3.92691* | .70415 | .000 | -6.2614 | -1.5924 |
| | F | -4.66022* | .70415 | .000 | -6.9948 | -2.3257 |
| | G | -5.66143* | .70415 | .000 | -7.9960 | -3.3269 |
| | H | -5.46252* | .70415 | .000 | -7.7970 | -3.1280 |
| | J | -5.85887* | .78726 | .000 | -8.4689 | -3.2488 |
| J | B | 2.16720 | .70415 | .086 | -.1673 | 4.5017 |
| | C | .80754 | .65867 | .945 | -1.3762 | 2.9913 |
| | D | 2.19555* | .60981 | .025 | .1738 | 4.2173 |
| | E | 1.93196 | .60981 | .070 | -.0898 | 3.9537 |
| | F | 1.19864 | .60981 | .576 | -.8231 | 3.2204 |
| | G | .19744 | .60981 | 1.000 | -1.8243 | 2.2192 |
| | H | .39635 | .60981 | .999 | -1.6254 | 2.4181 |
| | I | 5.85887* | .78726 | .000 | 3.2488 | 8.4689 |

*. The mean difference is significant at the 0.05 level.

**ANOVA (n=42)**

Nasal

| | Sum of Squares | df | Mean Square | F | Sig. |
|---|---|---|---|---|---|
| Between Groups | 21.088 | 8 | 2.636 | 2.321 | .043 |
| Within Groups | 37.483 | 33 | 1.136 | | |
| Total | 58.571 | 41 | | | |

This initial analysis reveals statistically significant differences between the groups as a whole. The table below displays the results of Multiple Comparisons between the individual groups (Tukey post hoc test).

**Multiple Comparisons**

For better reading, the infection doses were marked as follows: B: 1 x $10^5$ $TCID_{50}$, C: 1 x $10^3$ $TCID_{50}$, D: 1 x $10^2$ $TCID_{50}$, E: 1 x $10^1$ $TCID_{50}$, F: 1 x $10^0$ $TCID_{50}$, G: 1 x $10^{-1}$ $TCID_{50}$, H: 1 x $10^{-2}$ $TCID_{50}$, I: 1 x $10^{-3}$ $TCID_{50}$, J: 1 x $10^{-4}$ $TCID_{50}$. *P*-values below 0.05 were considered statistically significant and are marked in green.

| (I) Dose | (J) Dose | Mean Difference (I-J) | Std. Error | Sig. | 95% Confidence Interval | |
|---|---|---|---|---|---|---|
| | | | | | Lower Bound | Upper Bound |
| B | C | .78005 | .81399 | .987 | -1.9187 | 3.4788 |
| | D | -.39285 | .75361 | 1.000 | -2.8914 | 2.1057 |
| | E | .14956 | .75361 | 1.000 | -2.3490 | 2.6481 |
| | F | -.37987 | .75361 | 1.000 | -2.8784 | 2.1186 |
| | G | -.08403 | .75361 | 1.000 | -2.5825 | 2.4145 |
| | H | .03253 | .75361 | 1.000 | -2.4660 | 2.5310 |
| | I | 2.93828 | .97290 | .097 | -.2873 | 6.1639 |
| | J | .00719 | .87019 | 1.000 | -2.8778 | 2.8922 |
| C | B | -.78005 | .81399 | .987 | -3.4788 | 1.9187 |
| | D | -1.17290 | .68795 | .739 | -3.4537 | 1.1079 |
| | E | -.63048 | .68795 | .990 | -2.9113 | 1.6503 |
| | F | -1.15992 | .68795 | .750 | -3.4407 | 1.1209 |
| | G | -.86408 | .68795 | .937 | -3.1449 | 1.4167 |
| | H | -.74752 | .68795 | .972 | -3.0283 | 1.5333 |
| | I | 2.15823 | .92298 | .350 | -.9018 | 5.2183 |
| | J | -.77286 | .81399 | .988 | -3.4716 | 1.9258 |
| D | B | .39285 | .75361 | 1.000 | -2.1057 | 2.8914 |
| | C | 1.17290 | .68795 | .739 | -1.1079 | 3.4537 |
| | E | .54242 | .61532 | .993 | -1.4976 | 2.5824 |
| | F | .01298 | .61532 | 1.000 | -2.0270 | 2.0530 |
| | G | .30882 | .61532 | 1.000 | -1.7312 | 2.3489 |
| | H | .42538 | .61532 | .999 | -1.6146 | 2.4654 |
| | I | 3.33113* | .87019 | .014 | .4461 | 6.2162 |
| | J | .40004 | .75361 | 1.000 | -2.0985 | 2.8986 |
| E | B | -.14956 | .75361 | 1.000 | -2.6481 | 2.3490 |
| | C | .63048 | .68795 | .990 | -1.6503 | 2.9113 |
| | D | -.54242 | .61532 | .993 | -2.5824 | 1.4976 |
| | F | -.52943 | .61532 | .994 | -2.5695 | 1.5106 |
| | G | -.23359 | .61532 | 1.000 | -2.2736 | 1.8064 |
| | H | -.11703 | .61532 | 1.000 | -2.1571 | 1.9230 |
| | I | 2.78872 | .87019 | .065 | -.0963 | 5.6738 |
| | J | -.14237 | .75361 | 1.000 | -2.6409 | 2.3561 |
| F | B | .37987 | .75361 | 1.000 | -2.1186 | 2.8784 |
| | C | 1.15992 | .68795 | .750 | -1.1209 | 3.4407 |

| | | | | | | |
|---|---|---|---|---|---|---|
| | D | -.01298 | .61532 | 1.000 | -2.0530 | 2.0270 |
| | E | .52943 | .61532 | .994 | -1.5106 | 2.5695 |
| | G | .29584 | .61532 | 1.000 | -1.7442 | 2.3359 |
| | H | .41240 | .61532 | .999 | -1.6276 | 2.4524 |
| | I | 3.31815* | .87019 | .015 | .4331 | 6.2032 |
| | J | .38706 | .75361 | 1.000 | -2.1115 | 2.8856 |
| G | B | .08403 | .75361 | 1.000 | -2.4145 | 2.5825 |
| | C | .86408 | .68795 | .937 | -1.4167 | 3.1449 |
| | D | -.30882 | .61532 | 1.000 | -2.3489 | 1.7312 |
| | E | .23359 | .61532 | 1.000 | -1.8064 | 2.2736 |
| | F | -.29584 | .61532 | 1.000 | -2.3359 | 1.7442 |
| | H | .11656 | .61532 | 1.000 | -1.9235 | 2.1566 |
| | I | 3.02231* | .87019 | .034 | .1373 | 5.9073 |
| | J | .09122 | .75361 | 1.000 | -2.4073 | 2.5897 |
| H | B | -.03253 | .75361 | 1.000 | -2.5310 | 2.4660 |
| | C | .74752 | .68795 | .972 | -1.5333 | 3.0283 |
| | D | -.42538 | .61532 | .999 | -2.4654 | 1.6146 |
| | E | .11703 | .61532 | 1.000 | -1.9230 | 2.1571 |
| | F | -.41240 | .61532 | .999 | -2.4524 | 1.6276 |
| | G | -.11656 | .61532 | 1.000 | -2.1566 | 1.9235 |
| | I | 2.90575* | .87019 | .047 | .0207 | 5.7908 |
| | J | -.02534 | .75361 | 1.000 | -2.5239 | 2.4732 |
| I | B | -2.93828 | .97290 | .097 | -6.1639 | .2873 |
| | C | -2.15823 | .92298 | .350 | -5.2183 | .9018 |
| | D | -3.33113* | .87019 | .014 | -6.2162 | -.4461 |
| | E | -2.78872 | .87019 | .065 | -5.6738 | .0963 |
| | F | -3.31815* | .87019 | .015 | -6.2032 | -.4331 |
| | G | -3.02231* | .87019 | .034 | -5.9073 | -.1373 |
| | H | -2.90575* | .87019 | .047 | -5.7908 | -.0207 |
| | J | -2.93109 | .97290 | .099 | -6.1567 | .2945 |
| J | B | -.00719 | .87019 | 1.000 | -2.8922 | 2.8778 |
| | C | .77286 | .81399 | .988 | -1.9258 | 3.4716 |
| | D | -.40004 | .75361 | 1.000 | -2.8986 | 2.0985 |
| | E | .14237 | .75361 | 1.000 | -2.3561 | 2.6409 |
| | F | -.38706 | .75361 | 1.000 | -2.8856 | 2.1115 |
| | G | -.09122 | .75361 | 1.000 | -2.5897 | 2.4073 |
| | H | .02534 | .75361 | 1.000 | -2.4732 | 2.5239 |
| | I | 2.93109 | .97290 | .099 | -.2945 | 6.1567 |

*. The mean difference is significant at the 0.05 level.

**ANOVA (n=37)**

Trachea

| | Sum of Squares | df | Mean Square | F | Sig. |
|---|---|---|---|---|---|
| Between Groups | 63.994 | 8 | 7.999 | 3.601 | .005 |
| Within Groups | 62.205 | 28 | 2.222 | | |
| Total | 126.199 | 36 | | | |

This initial analysis reveals statistically significant differences between the groups as a whole. The table below displays the results of Multiple Comparisons between the individual groups (Tukey post hoc test).

**Multiple Comparisons**

For better reading, the infection doses were marked as follows: B: 1 x $10^5$ $TCID_{50}$, C: 1 x $10^3$ $TCID_{50}$, D: 1 x $10^2$ $TCID_{50}$, E: 1 x $10^1$ $TCID_{50}$, F: 1 x $10^0$ $TCID_{50}$, G: 1 x $10^{-1}$ $TCID_{50}$, H: 1 x $10^{-2}$ $TCID_{50}$, I: 1 x $10^{-3}$ $TCID_{50}$, J: 1 x $10^{-4}$ $TCID_{50}$. *P*-values below 0.05 were considered statistically significant and are marked in green.

| (I) Dose | (J) Dose | Mean Difference (I-J) | Std. Error | Sig. | 95% Confidence Interval | |
|---|---|---|---|---|---|---|
| | | | | | Lower Bound | Upper Bound |
| B | C | -.42501 | 1.29081 | 1.000 | -4.7556 | 3.9056 |
| | D | -.60660 | 1.21699 | 1.000 | -4.6896 | 3.4764 |
| | E | -.98117 | 1.36064 | .998 | -5.5461 | 3.5837 |
| | F | -3.31525 | 1.21699 | .185 | -7.3982 | .7677 |
| | G | -1.67049 | 1.24705 | .910 | -5.8543 | 2.5133 |
| | H | -2.14565 | 1.21699 | .704 | -6.2286 | 1.9373 |
| | I | -3.43573 | 1.49050 | .372 | -8.4363 | 1.5649 |
| | J | -4.36073 | 1.36064 | .070 | -8.9256 | .2042 |
| C | B | .42501 | 1.29081 | 1.000 | -3.9056 | 4.7556 |
| | D | -.18160 | .96212 | 1.000 | -3.4095 | 3.0463 |
| | E | -.55617 | 1.13839 | 1.000 | -4.3754 | 3.2631 |
| | F | -2.89024 | .96212 | .107 | -6.1181 | .3376 |
| | G | -1.24548 | .99986 | .939 | -4.6000 | 2.1090 |
| | H | -1.72065 | .96212 | .689 | -4.9485 | 1.5072 |
| | I | -3.01072 | 1.29081 | .358 | -7.3414 | 1.3199 |
| | J | -3.93572* | 1.13839 | .040 | -7.7550 | -.1165 |

| | | | | | | |
|---|---|---|---|---|---|---|
| D | B | .60660 | 1.21699 | 1.000 | -3.4764 | 4.6896 |
| | C | .18160 | .96212 | 1.000 | -3.0463 | 3.4095 |
| | E | -.37457 | 1.05395 | 1.000 | -3.9105 | 3.1614 |
| | F | -2.70865 | .86054 | .079 | -5.5957 | .1784 |
| | G | -1.06389 | .90255 | .955 | -4.0919 | 1.9641 |
| | H | -1.53905 | .86054 | .689 | -4.4261 | 1.3480 |
| | I | -2.82913 | 1.21699 | .362 | -6.9121 | 1.2538 |
| | J | -3.75413* | 1.05395 | .031 | -7.2901 | -.2182 |
| E | B | .98117 | 1.36064 | .998 | -3.5837 | 5.5461 |
| | C | .55617 | 1.13839 | 1.000 | -3.2631 | 4.3754 |
| | D | .37457 | 1.05395 | 1.000 | -3.1614 | 3.9105 |
| | F | -2.33408 | 1.05395 | .424 | -5.8700 | 1.2019 |
| | G | -.68932 | 1.08851 | .999 | -4.3412 | 2.9626 |
| | H | -1.16448 | 1.05395 | .969 | -4.7004 | 2.3715 |
| | I | -2.45456 | 1.36064 | .679 | -7.0194 | 2.1103 |
| | J | -3.37956 | 1.21699 | .167 | -7.4625 | .7034 |
| F | B | 3.31525 | 1.21699 | .185 | -.7677 | 7.3982 |
| | C | 2.89024 | .96212 | .107 | -.3376 | 6.1181 |
| | D | 2.70865 | .86054 | .079 | -.1784 | 5.5957 |
| | E | 2.33408 | 1.05395 | .424 | -1.2019 | 5.8700 |
| | G | 1.64476 | .90255 | .668 | -1.3832 | 4.6728 |
| | H | 1.16959 | .86054 | .904 | -1.7175 | 4.0567 |
| | I | -.12048 | 1.21699 | 1.000 | -4.2034 | 3.9625 |
| | J | -1.04548 | 1.05395 | .984 | -4.5814 | 2.4905 |
| G | B | 1.67049 | 1.24705 | .910 | -2.5133 | 5.8543 |
| | C | 1.24548 | .99986 | .939 | -2.1090 | 4.6000 |
| | D | 1.06389 | .90255 | .955 | -1.9641 | 4.0919 |
| | E | .68932 | 1.08851 | .999 | -2.9626 | 4.3412 |
| | F | -1.64476 | .90255 | .668 | -4.6728 | 1.3832 |
| | H | -.47516 | .90255 | 1.000 | -3.5032 | 2.5528 |
| | I | -1.76524 | 1.24705 | .882 | -5.9490 | 2.4186 |
| | J | -2.69024 | 1.08851 | .287 | -6.3421 | .9617 |
| H | B | 2.14565 | 1.21699 | .704 | -1.9373 | 6.2286 |
| | C | 1.72065 | .96212 | .689 | -1.5072 | 4.9485 |
| | D | 1.53905 | .86054 | .689 | -1.3480 | 4.4261 |
| | E | 1.16448 | 1.05395 | .969 | -2.3715 | 4.7004 |
| | F | -1.16959 | .86054 | .904 | -4.0567 | 1.7175 |
| | G | .47516 | .90255 | 1.000 | -2.5528 | 3.5032 |
| | I | -1.29007 | 1.21699 | .975 | -5.3730 | 2.7929 |
| | J | -2.21508 | 1.05395 | .492 | -5.7510 | 1.3209 |

| | | | | | | |
|---|---|---|---|---|---|---|
| I | B | 3.43573 | 1.49050 | .372 | -1.5649 | 8.4363 |
| | C | 3.01072 | 1.29081 | .358 | -1.3199 | 7.3414 |
| | D | 2.82913 | 1.21699 | .362 | -1.2538 | 6.9121 |
| | E | 2.45456 | 1.36064 | .679 | -2.1103 | 7.0194 |
| | F | .12048 | 1.21699 | 1.000 | -3.9625 | 4.2034 |
| | G | 1.76524 | 1.24705 | .882 | -2.4186 | 5.9490 |
| | H | 1.29007 | 1.21699 | .975 | -2.7929 | 5.3730 |
| | J | -.92500 | 1.36064 | .999 | -5.4899 | 3.6399 |
| J | B | 4.36073 | 1.36064 | .070 | -.2042 | 8.9256 |
| | C | 3.93572* | 1.13839 | .040 | .1165 | 7.7550 |
| | D | 3.75413* | 1.05395 | .031 | .2182 | 7.2901 |
| | E | 3.37956 | 1.21699 | .167 | -.7034 | 7.4625 |
| | F | 1.04548 | 1.05395 | .984 | -2.4905 | 4.5814 |
| | G | 2.69024 | 1.08851 | .287 | -.9617 | 6.3421 |
| | H | 2.21508 | 1.05395 | .492 | -1.3209 | 5.7510 |
| | I | .92500 | 1.36064 | .999 | -3.6399 | 5.4899 |

*. The mean difference is significant at the 0.05 level.
